# Inferring and Evaluating Network Medicine-Based Disease Modules with Nextflow

**DOI:** 10.1101/2025.11.20.687681

**Authors:** Johannes Kersting, Chloé Bucheron, Lisa M. Spindler, Joaquim Aguirre-Plans, Quirin Manz, Tanja Pock, Mo Tan, Fernando M. Delgado-Chaves, Cristian Nogales, Harald H. H. W. Schmidt, Jörg Menche, Andreas Maier, Jan Baumbach, Emre Guney, Markus List

**Affiliations:** Data Science in Systems Biology, TUM School of Life Sciences, Technical University of Munich, Freising, Germany; Max Perutz Labs, Department of Structural and Computational Biology, University of Vienna, Vienna, Austria; Vienna Biocenter PhD Program, a Doctoral School of the University of Vienna and the Medical University of Vienna, Vienna, Austria; Ludwig Boltzmann Institute for Network Medicine at the University of Vienna, Vienna, Austria; Discovery and Data Science (DDS) Unit, STALICLA SL, Barcelona, Spain; Institute for Computational Systems Biomedicine, University of Hamburg, Hamburg, Germany; Department of Pharmacology and Personalised Medicine, Maastricht University, Maastricht, The Netherlands; Faculty of Mathematics, University of Vienna, Vienna, Austria; CeMM Research Center for Molecular Medicine of the Austrian Academy of Sciences, Vienna, Austria; Munich Data Science Institute (MDSI), Technical University of Munich, Garching, Germany

**Keywords:** network medicine, disease modules, drug repurposing, pipeline, Nextflow

## Abstract

Most human diseases result from complex molecular interactions of genes and proteins. Various network-based computational methods characterize these mechanisms by expanding seed genes into disease-associated subnetworks, or disease modules.

Evaluating these diverse methods is tedious due to unique installation and data preparation requirements. Moreover, the underlying algorithmic strategies differ, making it difficult to determine which of the created modules are most useful or biologically plausible.

To address this challenge, we developed an all-in-one Nextflow pipeline that enables automated and reproducible analyses. It handles installation, input preparation, execution, and systematic evaluation of six widely used module detection tools, considering module topology, functional coherence, robustness, and the capacity to recover seeds. In addition, it annotates the resulting disease modules with biological context information, prioritizes potential drug candidates, and generates visualizations and a comprehensive summary report.

To showcase the value of our pipeline and offer guidance to potential users, we performed a comprehensive evaluation across 50 different disease-network combinations, revealing substantial variability among the derived disease modules. We show that this variability is driven by differences in modeling approach, input network, and seed composition. While most methods are robust to minor perturbations, they struggle to recover omitted seeds, and none consistently outperforms others, underscoring the need for careful method selection.

Our work enables the research community to systematically compare approaches for disease module discovery, promoting reproducible network medicine research. Integrated into the nf-core project (https://nf-co.re/diseasemodulediscovery), it is intended as an extendable, long-term resource for tracking progress in the field.

## Introduction

Diseases are seldom purely monogenic but often emerge from complex disturbances in the interactions between different genes and biomolecules. These molecular interactions collectively form the human interactome, which can be represented as a network in which nodes correspond to biomolecules, i.e., proteins, DNA, or metabolites, and edges model the interactions between them. Studying the interactome and its disease-related perturbations is a central area in network medicine [1]. A key insight of the field is that disease-associated genes, or more precisely, their protein products, are not randomly scattered throughout the interactome. Instead, they tend to cluster within local neighborhoods [2], forming so-called disease modules: localized subnetworks of the interactome encompassing disease-related proteins and their interactions [3,4].

Disease modules thus characterize diseases through their molecular mechanisms, rather than traditional organ- or symptom-based definitions, implementing the foundational idea of systems medicine. Expanding this view towards network pharmacology shows therapeutic potential, as disease modules can be used to discover novel drug targets and drug candidates that precisely address the underlying pathological processes. Targeting multiple components within a disease module may further enhance efficacy through synergistic drug effects [5]. Moreover, this approach synergizes with drug repurposing, i.e., using approved drugs with known protein targets for new indications. Compared to traditional drug development, repurposing can significantly reduce time, cost, and risk [6]. Network medicine and disease modules have been demonstrated to be effective strategies for drug repurposing [7], and these concepts have been applied to study a wide range of diseases, including cancer [7], SARS-CoV-2 infection [8], hypertension [9], and stroke [10].

Given the potential of disease modules to provide mechanistic insights into diseases and support drug repurposing, numerous algorithmic approaches for their identification have been proposed. A common approach involves starting with a set of known disease-associated genes or proteins (referred to as seeds) and expanding them into disease modules by incorporating additional nodes from the interactome (see Figure 1A). The inclusion of nodes is guided by their connectivity and proximity to the seed set within the network, implicating them as key players in the disease mechanism.

**Figure 1:**
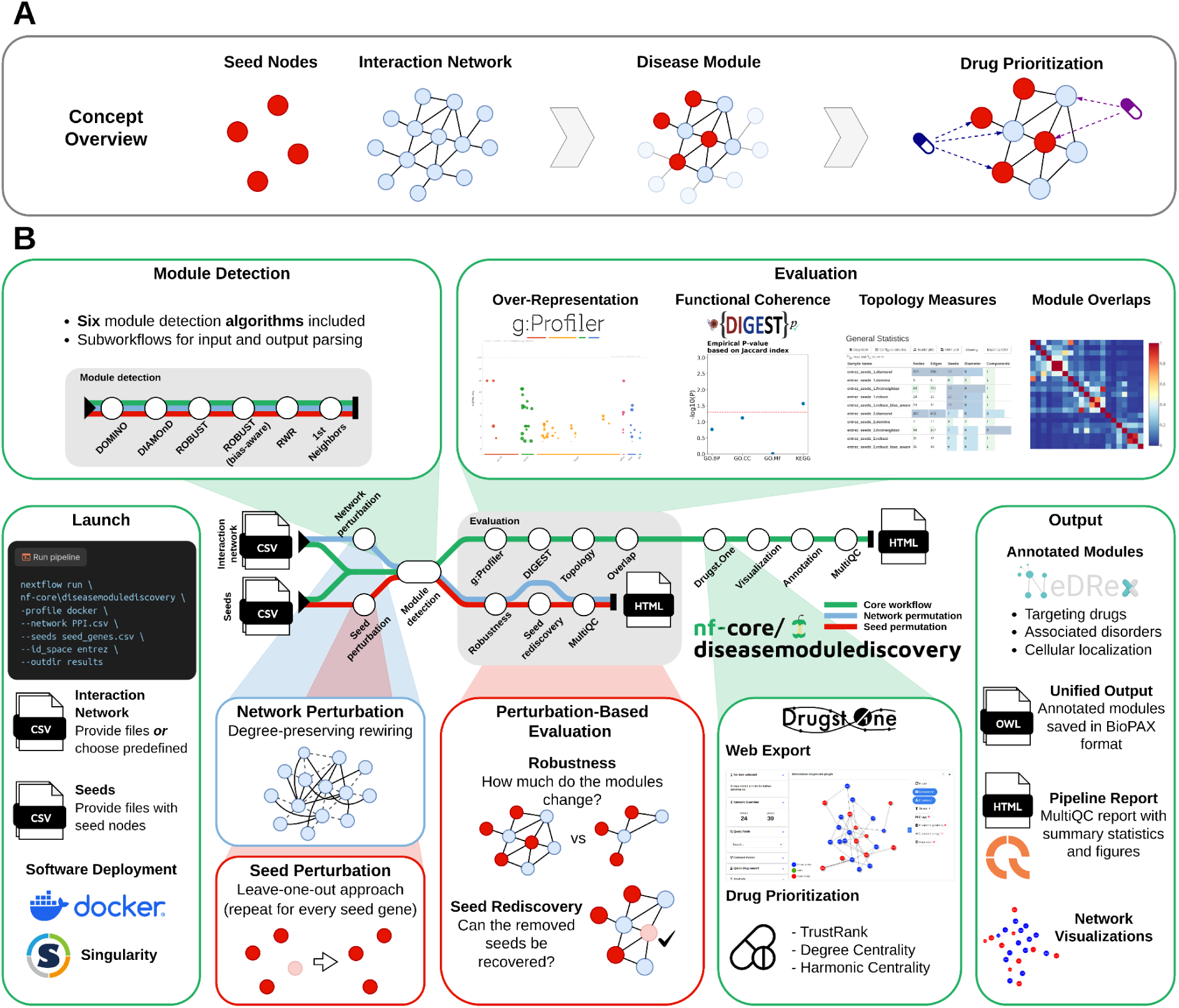
(**A**) Overview of a network medicine–based drug repurposing workflow, starting from the seed set and input network, through disease module identification, to drug prioritization. (**B**) Our pipeline infers disease modules using six widely used approaches. The resulting modules are evaluated through overrepresentation and functional coherence analyses, network topology measures, and pairwise node set overlaps, enabling comparison of modules derived from different seed sets, networks, or AMIMs. Degree-preserving network rewiring is applied to assess potential node-degree biases. A leave-one-out analysis on the seed set evaluates the robustness of the AMIMs to small input perturbations and calculates a rediscovery rate. Drug candidates are then prioritized using various network-based algorithms. The pipeline also generates network visualizations of the modules, annotates them with supplementary biological information, and provides an HTML summary report of the entire run.

A dilemma of disease module discovery is the non-trivial method choice. Most disease module discovery methods, also referred to as active module identification methods (AMIMs), differ not only by their algorithmic approach but also by following a different rationale on how a disease module is best identified. A First-Nearest Neighbors (1st Neighbors) approach, for example, simply expands the seed set by adding all nodes that directly interact with at least one seed node. DIAMOnD [2] iteratively expands the seed nodes by repeatedly adding the node with the most significant number of interactions with the current seeds, considering its overall node degree. DOMINO [11] first partitions the whole network into non-overlapping clusters (slices), then returns and further refines those with a higher-than-expected number of seeds. ROBUST [12] repeatedly connects the seed nodes into a single connected component by building Prize-Collecting Steiner Trees [13]. Nodes that appear in a sufficiently large number of trees are incorporated into the final solution. Its research bias-aware implementation [14] works similarly, but penalizes the inclusion of heavily overstudied proteins by considering the number of times a protein was used as bait in protein-protein interaction (PPI) detection experiments. Network propagation methods, such as random walk with restart (RWR) [15], can be used to simultaneously expand the seed set in all directions, e.g., until they constitute a single connected component.

This diversity of methods is required, as the best approach may depend on both the topological shape of the disease mechanism (e.g., organized around a single hub, describing a pathway, or more complex patterns) but also on the abundance of and confidence in known seed genes. For example, if a disease mechanism is characterized by multiple disjoint subnetworks, a method such as ROBUST, which is based on connecting the seeds into a single component, is inappropriate. If only a few disease-associated genes are known, a more explorative method may yield better results than a more conservative method. For most disease mechanisms, we can only speculate on these properties, necessitating a framework that allows us to assess multiple approaches in parallel and to compare them effectively.

Such an assessment of disease modules can focus on overlaps with known pathways [2,11,12,16] or by analyzing their functional coherence [17]. Another common evaluation strategy is to analyze the robustness of the methods and their outputs under various types of input perturbations [2,11,16]. However, applying and comparing several AMIMs is hindered by method-specific installation processes, input requirements, and output formats, making manual execution cumbersome. Moreover, input perturbation-based evaluations are computationally demanding and necessitate efficient parallelization to remain practical. When further combined with downstream analyses such as visualization and drug candidate prioritization, disease module discovery thus becomes a complex, multi-step workflow that requires a diverse set of tools.

Several web-based platforms [7,8,18–20] improve accessibility by offering unified access to various AMIMs, network visualizations, and drug prioritization algorithms without requiring installation. However, they are limited in terms of automation, scalability, and the range of evaluation strategies they support. Lazareva et al. [16] introduced an automated test suite to benchmark AMIMs, but it is designed for use with predefined datasets only and is limited to assessing the impact of network perturbations. Yang et al. [21] conducted a benchmarking analysis using synthetic and mutation data, which is also limited to predefined benchmarking datasets and specifically tailored toward cancer applications. Disease module identification has also been evaluated through a DREAM challenge [22]. However, methods were restricted to inferring modules solely from network structure, without incorporating disease-specific context through seed genes.

As part of the Horizon Europe project REPO4EU (https://repo4.eu), which aims to develop a comprehensive platform for drug repurposing, we sought a robust workflow that includes disease module identification, systematic evaluation, visualization, and drug prioritization. To close this gap, we developed a Nextflow [23] pipeline that autonomously executes all these steps, parallelizes tasks where possible, and manages dependencies via containers, simplifying the setup and ensuring reproducibility. The pipeline is released as part of the nf-core framework [24], a collection of community-curated bioinformatics workflows. To showcase the analytical power of our pipeline, we applied it to a combination of ten diverse input networks and five seed sets. In addition to demonstrating the volume and diversity of results that can be generated from a single pipeline run, it also enabled us to make several key observations about disease modules and their corresponding AMIMs: (i) reflecting their distinct modeling approaches, different AMIMs produce disease modules with markedly different topologies; (ii) most AMIMs produce disease modules that are specific to their corresponding diseases, although this specificity largely arises from the shared seed set; (iii) the choice of input network has a major influence on the resulting modules; (iv) most AMIMs exhibit robustness to small input perturbations; but (v) generally struggle to reliably rediscover individually omitted seeds; (vi) no AMIM consistently outperformed the others across all categories, underscoring the need for careful method selection.

## Results

### A pipeline for reproducible disease module identification and drug repurposing

Our pipeline (see Figure 1B) accepts one or more seed files and one or more networks as input. Seed files must be provided by the user and can come from disease databases such as DisGeNet [25,26], ClinVar [27], or Open Targets [28], genome-wide association studies (GWAS), literature searches, or own research results, e.g., from differential expression or genomic variant calling analyses. Networks can either be supplied or selected from a list of available options (see Table S1).

The pipeline then runs six widely used algorithms to infer disease modules based on the provided input. This includes DOMINO [11], DIAMOnD [2], ROBUST [12] and its bias-aware implementation [14], a 1st neighbors approach [9], and random walk with restart (RWR) [15]. By default, all methods are applied, but each can be easily excluded if desired. This part of the pipeline generates one module for every combination of seed file, network file, and AMIM specified. Each is then processed further individually.

Since disease modules usually lack a ground truth, in silico validation is challenging but essential for assessing the reliability of the results. Therefore, the pipeline includes extensive evaluation steps.

To get a general idea of the topology of the inferred modules, measures such as the number of nodes and edges, the number of included seeds, the diameter, the number of connected components, the size of the largest connected component (LCC), and the maximum distance from an added node to its nearest seed are reported (see Methods).

A common strategy for assessing the biological relevance of inferred modules is to test for enrichment in gene sets, such as biological pathways, known disease genes, or Gene Ontology (GO) terms [2,11,12]. This is typically done through over-representation analysis, which identifies gene sets linked to specific functions and significantly enriched with the genes in a module. The results can then be analyzed in the context of the disease of interest, providing insights into the biological processes captured by the module and its reliability by comparing the findings to user expectations. To support this analysis, the pipeline runs g:Profiler [29] to report enriched gene sets for each disease module. It also integrates DIGEST [17], a tool specifically developed to evaluate network modules by testing their functional coherence based on the premise that genes within a module should be associated with similar gene sets.

In addition to evaluating biological relevance, disease modules and their inference methods can be assessed by perturbing the input data. By removing seeds from the original set, it is possible to evaluate the robustness and seed rediscovery potential [2]. Specifically, the pipeline performs a leave-one-out analysis by removing one seed at a time and re-running the AMIMs for each modified input. The resulting modules are then compared to the original to evaluate their stability in response to small changes and to determine the extent to which they depend on individual seeds. This procedure also enables calculating a seed rediscovery rate [30], i.e., the probability that a left-out seed is re-included in the inferred module. This metric estimates the AMIM’s ability to recover disease-associated genes or proteins not initially present in the input. Conversely, an overall low rediscovery rate across methods may hint at a low-confidence seed.

Previous research has shown that many AMIMs rely heavily on node degrees in the input network rather than on the information carried by individual edges [16]. To detect this bias, the pipeline repeatedly perturbs the input network by rewiring its edges while preserving node degrees. It then applies the AMIMs to these perturbed networks. If the resulting modules remain largely unchanged, the AMIM exploits node degree information rather than specific interactions.

To support drug repurposing hypothesis generation, the pipeline prioritizes candidate drugs for each inferred module using network-based algorithms that rank compounds based on their connectivity or proximity to the disease module. This is enabled by interfacing with Drugst.One [18], a web tool for drug repurposing and interactive network visualization, via API calls.

To manually inspect the modules and prioritized drugs, the pipeline saves them in multiple output formats, including visual graph representations. In addition to static figures, this includes HTML files, allowing users to drag and rearrange nodes and export links to visualize and manipulate the modules directly in Drugst.One.

Additionally, the pipeline annotates the modules with supplementary biological information. These data are retrieved from the NeDRex database [31] and include associated disorders, subcellular localization of proteins, as well as targeting drugs, their side effects, indications, and contraindications. To represent the annotated modules in a standardized manner, the Biological Pathway Exchange (BioPAX) [32] Level 3 format is used.

Finally, a summary report for the pipeline is generated using MultiQC [33]. This report provides an overview of the run, summarizing the topological properties of the modules, their overlaps, and the evaluation results from both DIGEST and the perturbation-based analyses, all in one place.

The pipeline code and documentation are openly available through GitHub (https://github.com/nf-core/diseasemodulediscovery) and the nf-core website (https://nf-co.re/diseasemodulediscovery).

### Lessons learned from applying systematic disease module discovery to a broad set of diseases and networks

To showcase the pipeline’s functionality and highlight characteristics of the integrated AMIMs, we applied it to five different seed sets combined with ten PPI networks, yielding 50 input combinations. The seed sets consisted of disease-associated genes for Huntington’s disease (HD), Ulcerative colitis (UC), Crohn’s disease (CD), Amyotrophic lateral sclerosis (ALS), and Lung adenocarcinoma (LUAD), obtained from DisGeNET (https://www.disgenet.com/) [25,26] (see Figure 2B, Table S2, and Methods for details). This selection of diseases was inspired by previous benchmarking approaches [12,14,16]. As interaction networks, we used human PPIs from STRING [34], BioGRID [35], HIPPIE [36], IID [37], and NeDRex [31] in different subset configurations corresponding to varying confidence levels of the interactions (see Figure 2A, Table S1, and Methods for details). All networks used in this demonstration are accessible through the pipeline interface, without the need to provide external input files.

**Figure 2:**
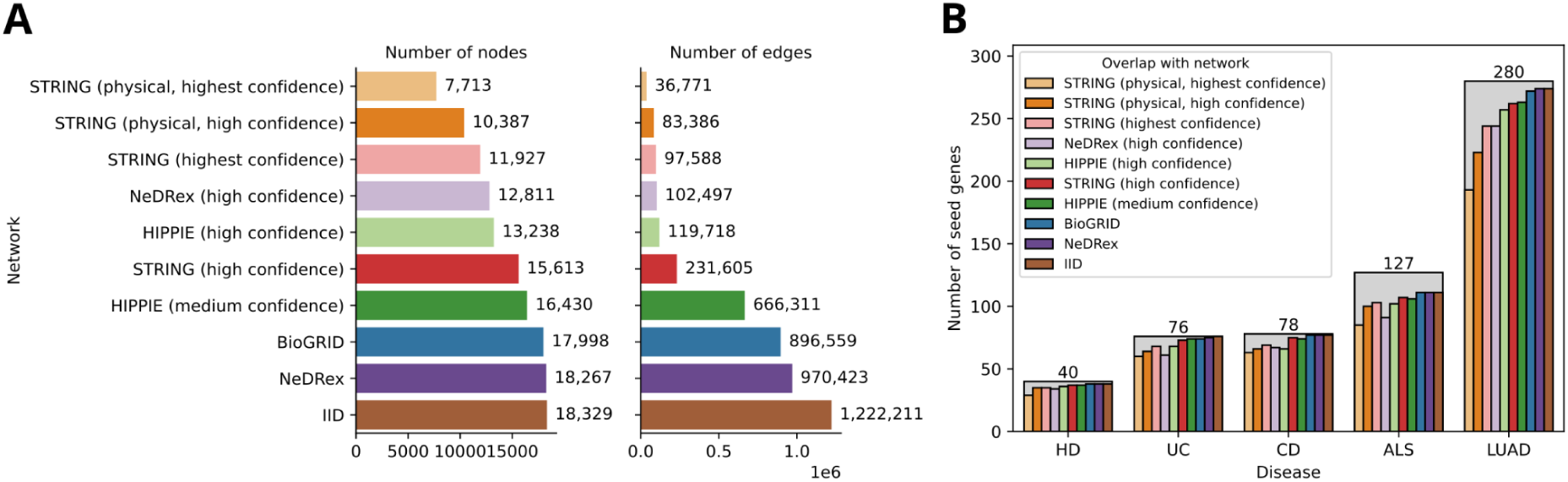
Inputs for the pipeline demonstration. (**A**) Number of nodes and edges across the ten input PPI networks. (**B**) Size of the five disease gene sets, shown as the number of seed nodes (from DisGeNET) (gray bars with annotations), and their overlaps with the node sets of the input networks (colored bars).

All analysis results presented in the following sections were generated using a single pipeline run. The raw outputs are available on Zenodo (https://zenodo.org/records/17536307). The “Only seeds” condition refers to the unmodified seed sets, without the inclusion of any additional nodes.

### AMIMs produce modules with distinct topologies

As expected, the AMIMs in the pipeline generated disease modules that differ regarding topological features such as module size, the number of connected components, the proportion of retained seeds, and their explorativeness. All measures were calculated on the subnetwork induced by the disease module (see Methods). It is important to note that some of the described properties may vary significantly depending on the specific parameterization of the AMIM (see Discussion).

DOMINO produces the smallest disease modules, followed by ROBUST, ROBUST (bias-aware), and DIAMOnD (Figure 1A). In contrast, RWR and 1st Neighbors can yield considerably larger modules, which can include thousands, and in the case of 1st Neighbors, even over ten thousand nodes. Naturally, module size is also influenced by the input data: all methods tend to produce larger modules when provided with larger seed sets (Figure S1). However, the effect of input network size varies by method (Figure 1E). In larger networks, seed nodes typically have more direct interactors, which leads to larger modules for the 1st Neighbor approach. Conversely, methods designed to connect seed nodes, such as ROBUST and RWR, generate smaller modules in larger networks, as additional nodes and edges allow for more efficient paths between seeds. The size of the modules produced by DOMINO, DIAMOnD, and ROBUST (bias-aware) is not significantly influenced by network size. This is notable in the case of ROBUST (bias-aware), since it also aims to connect the seeds, which can be done more efficiently in larger networks. A likely explanation is that ROBUST (bias-aware) penalizes the inclusion of highly connected, overstudied hub genes, thereby avoiding shortcuts in large networks that might otherwise reduce module size.

Interestingly, the seed nodes themselves never form a single connected component (Figure 1B). This may be due to the incompleteness of the interactome, the involvement of multiple independent mechanisms, or the existence of nodes relevant to the disease mechanism that are not yet included in the seed set, motivating the use of disease module inference algorithms to complete them [38]. The modules produced by DIAMOnD often consist of many components, but naturally fewer than the seeds form alone. DOMINO, ROBUST, and 1st Neighbors can yield multiple connected components, but generally return fewer than DIAMOnD. ROBUST (bias-aware) almost always returns a single connected component, while RWR always does so by design.

Except for DIAMOnD and 1st Neighbors, the evaluated AMIMs do not necessarily retain all input seed nodes in the resulting modules (Figure 1C). ROBUST, ROBUST (bias-aware), and RWR exclude seeds only when these are not part of the LCC of the input network. In contrast, DOMINO frequently omits seeds, which is expected given its approach, with a median retention rate of less than 50%.

Figure 1D illustrates the explorativeness of the AMIMs, defined as the frequency with which they add nodes that are distant from the seed nodes. As 1st Neighbors only includes direct interactors, this distance is always exactly one. Notably, DOMINO tends to incorporate primarily direct interactors, although no such restriction is built into the method. RWR and ROBUST also frequently return modules composed mainly of direct interactors, but second neighbors are commonly included as well. DIAMOnD typically incorporates at least second neighbors and occasionally even third neighbors. ROBUST (bias-aware) is by far the most explorative method, including nodes up to six edges away from any seed. This behavior is likely explained by the penalty imposed on overstudied hub genes [39], which would otherwise act as shortcuts connecting the seeds. Notably, the modules of ROBUST and ROBUST (bias-aware) can contain added nodes that are completely isolated from the seed nodes in the final module.

The observation that the modules generated by different AMIMs exhibit distinct topologies is consistent with observations from the module identification DREAM challenge [22]. It underscores the value of applying and comparing multiple methods for a single use case, as their solutions may be complementary.

### The included seed nodes are the primary reason for disease-specific modules

Despite the pronounced size differences between the input networks (Figure 3), we expect disease modules to be disease-specific regardless of the chosen network. Consequently, modules associated with the same disease should cluster together based on their similarity.

**Figure 3:**
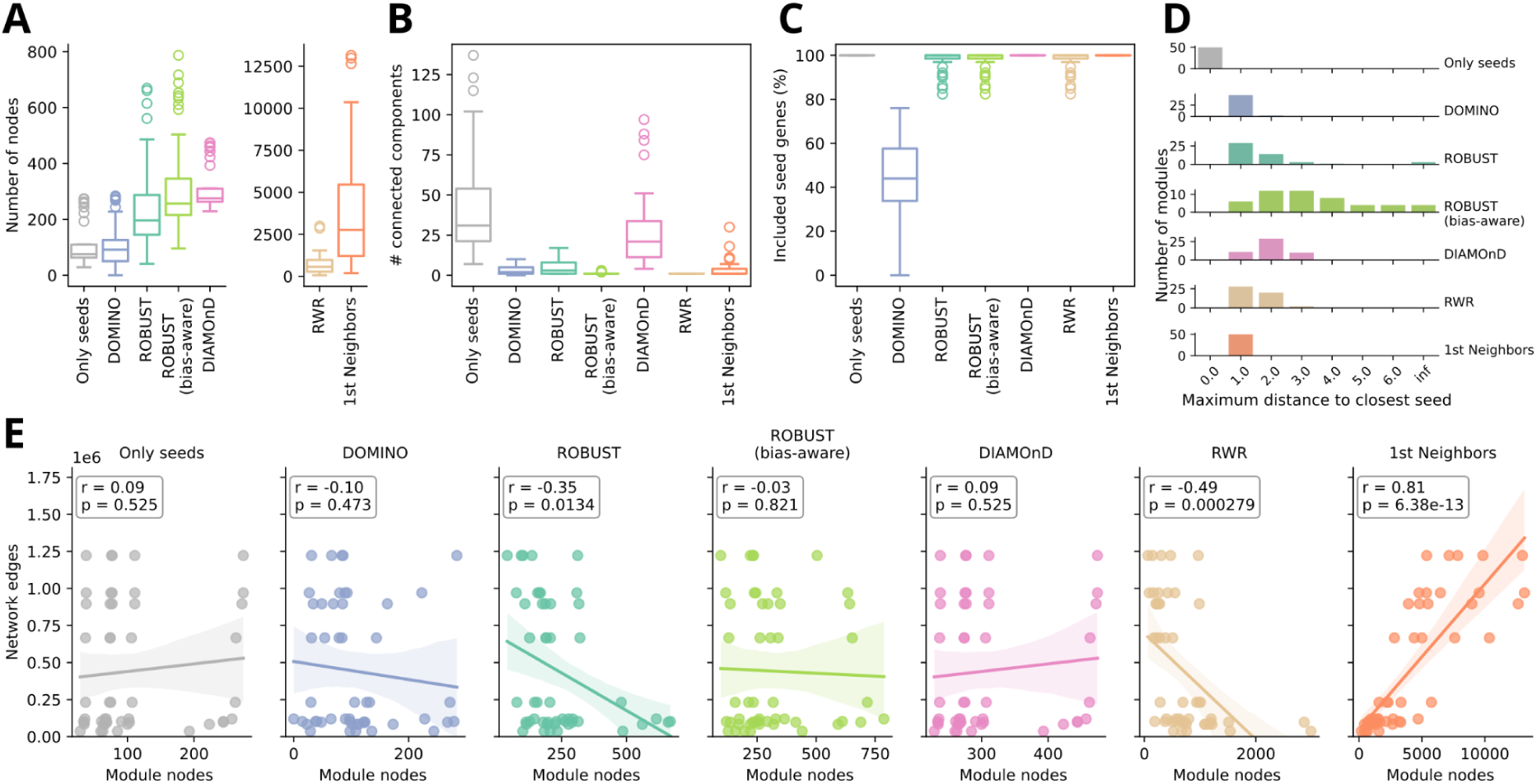
Topological properties of disease modules inferred using different methods. Subfigures **A**-**D** provide summaries across all 50 input combinations used. (**A**) Disease module size, defined as the number of nodes included in the module. (**B**) Number of connected components within each disease module. (**C**) Proportion of seed nodes retained in the final disease module. (**D**) Maximum distance from any added (non-seed) node to its nearest seed node, serving as a measure of explorativeness. A distance of 0 indicates that no additional nodes were included; “inf” (infinite) refers to nodes completely isolated from the seed set. (**E**) Correlation between disease module size and the number of edges in the input network. *r* indicates the Pearson correlation and *p* the corresponding p-value.

Figure 4A (see Figure S2 for a combined heatmap) confirms this expectation for modules generated by DOMINO, ROBUST, and ROBUST (bias-aware). Additionally, clear similarities can be observed between the modules for UC and CD, which is consistent with the fact that both are subtypes of Inflammatory Bowel Disease (IBD) and share mechanistic features through overlapping pathways [40]. In contrast, modules produced by DIAMOnD, RWR, and 1st Neighbors fail to cluster cleanly by disease. Since these methods generally generate larger modules (Figure 3A), this loss of specificity is likely caused by the inclusion of too many additional nodes.

**Figure 4:**
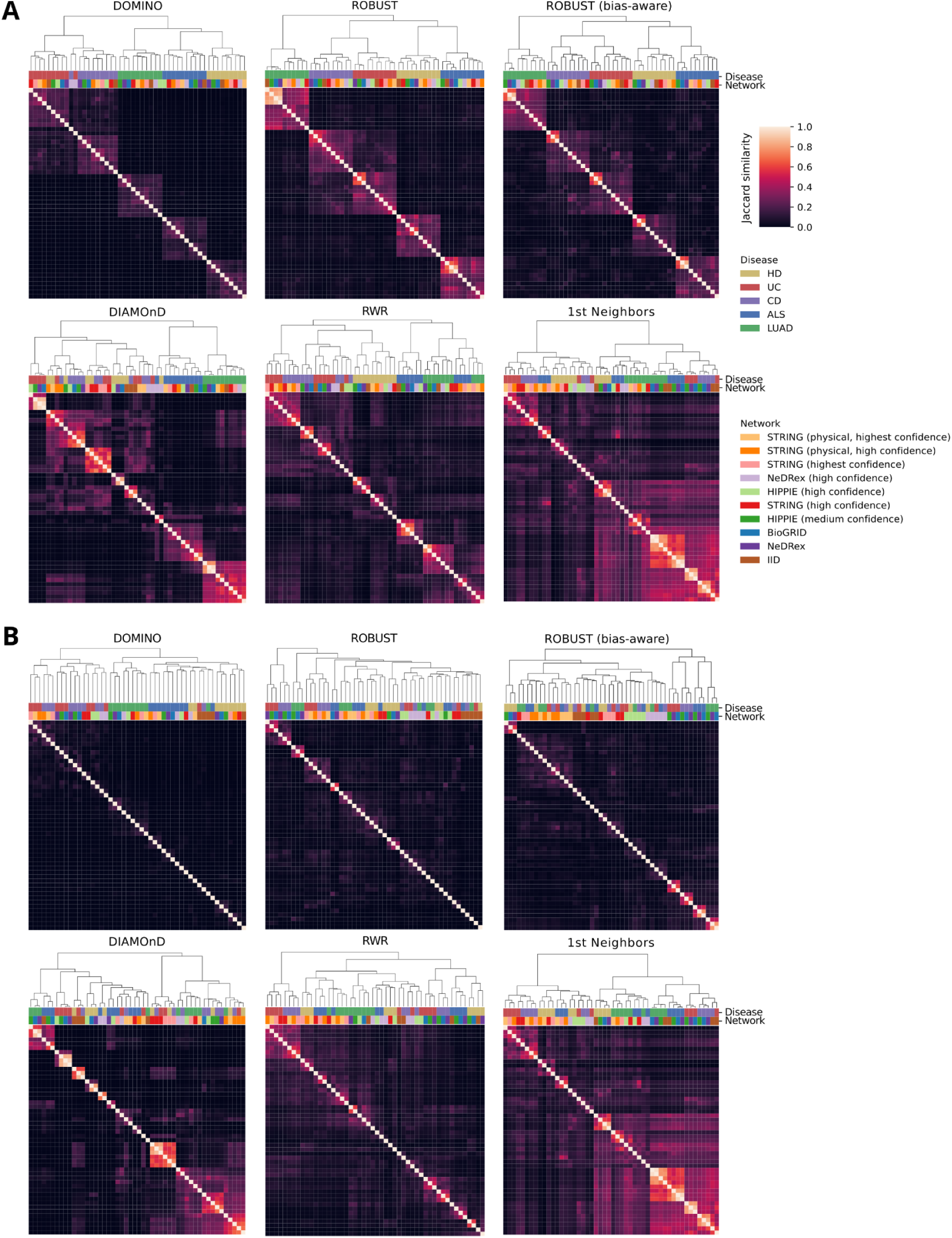
Heatmaps and hierarchical clusterings for each AMIM based on the pair-wise node set similarities of modules inferred using different diseases and networks. (**A**) Considering all module nodes. (**B**) Only considering added nodes (no seed nodes).

However, achieving disease-specific modules is a relatively low bar because the seed sets are already disease-specific and are typically retained in the resulting modules (Figure 3C). To address this, the pipeline also calculates module similarities while excluding the seed nodes and considering only the added nodes. Under this criterion (see Figure 4B and Figure S3 for a combined heatmap), the modules of most AMIMs no longer cluster by disease, indicating that the observed disease-specificity of the full modules mainly arises from the shared seed sets, rather than from the disease-specific added nodes. The only exception is DOMINO, which shows difficulty distinguishing the two IBD subtypes, UC and CD. Nevertheless, it largely maintains disease specificity, albeit with rather low overlaps (<0.25) off the diagonal.

### Disease modules are heavily influenced by network choice

As the results of the previous section indicated, the inferred modules are strongly affected by the choice of the input network. Figure 5 shows that the median Jaccard similarities between modules generated with the same AMIM and seed set, but different networks, are consistently below 0.5, even when the seed nodes are included in the overlap calculation. When only the added nodes are considered, the medians drop even further (<0.1 for DOMINO, ROBUST, and ROBUST (bias-aware); <0.2 for DIAMOnD and RWR; <0.4 for 1st Neighbors).

**Figure 5:**
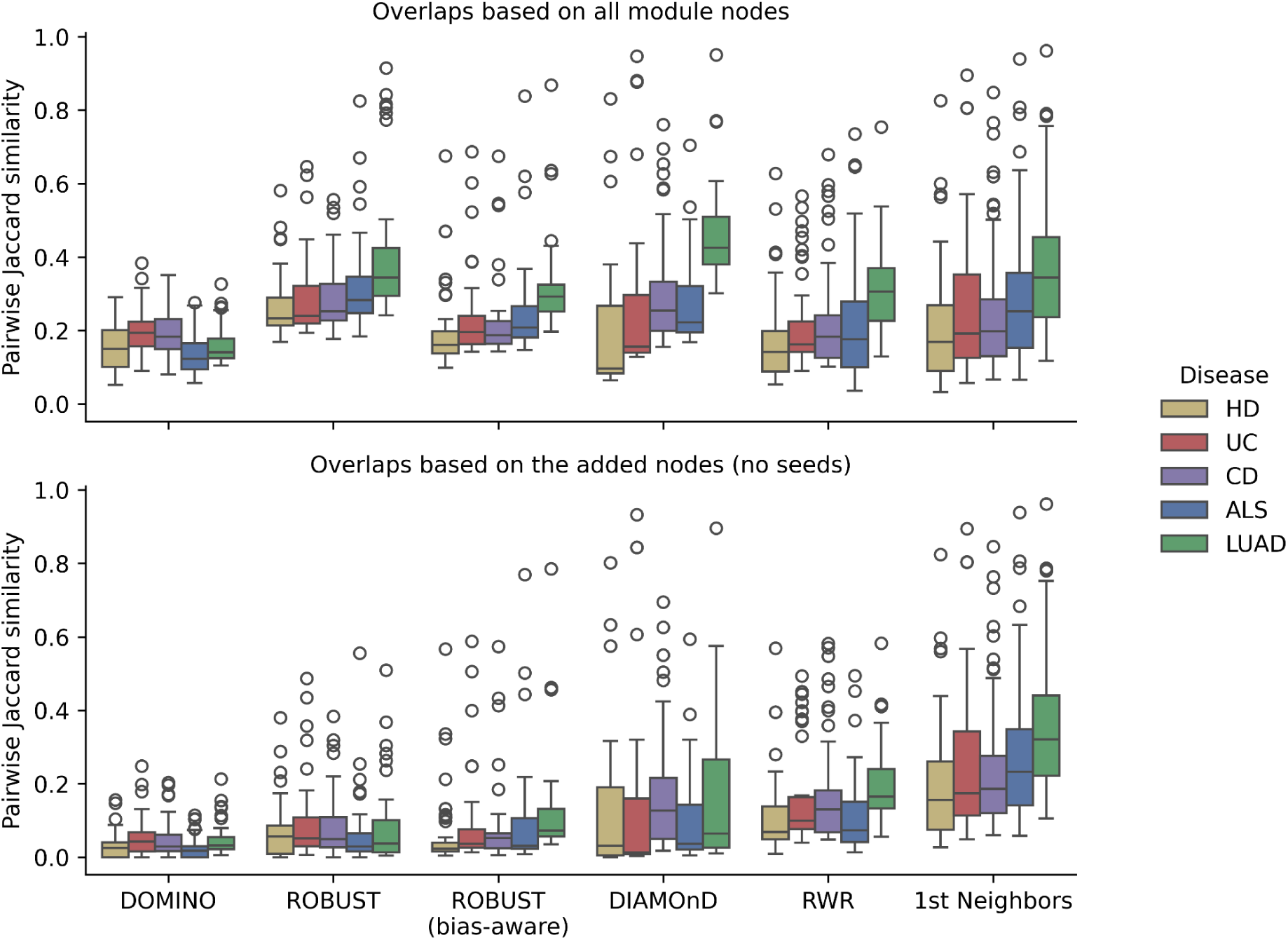
Distributions of the pair-wise Jaccard similarities between the node sets of disease modules inferred using the same seed set and AMIM, but different input networks. Top: considering all module nodes. Bottom: only considering added nodes (no seed nodes).

When all module nodes are considered, ROBUST produces the most consistent results across different networks, which aligns with the findings of Buzzao et al. [38]. However, when only the added nodes are considered, DIAMOnD, RWR, and 1st Neighbors exhibit higher consistency, despite their lower specificity in the previous analysis.

Overall, the low overlaps between modules inferred using different networks indicate that they are highly dependent on the chosen network. Consequently, a module inferred using one network is unlikely to be reproducible with a different network.

### Most AMIMs are robust to leave-one-out perturbations

Because gene–disease association data inevitably contain noise and false positives, disease modules should not rely excessively on individual seed nodes [2]. To evaluate this, the pipeline performs a leave-one-out analysis, sequentially removing each seed node from the input and rerunning the AMIMs to assess their robustness to small perturbations in the input by comparing the perturbed modules to the original ones using Jaccard similarity.

Figure 6A demonstrates that most AMIMs are robust to the omission of individual genes, with median mean Jaccard similarities exceeding 0.9. The only notable exception is DOMINO, which exhibits lower robustness (median ∼0.75). Notably, a clear positive relationship exists between module size and robustness (Figure 6B and Figure S4). Consequently, AMIMs that generate larger modules (Figure 3A) tend to yield more robust results. Moreover, with the exception of DOMINO, AMIMs also show increased robustness when provided with larger seed sets (Figure 6C), which is expected, as the relative contribution of any single seed decreases with larger seed sets. In contrast, the number of edges in the input network is negatively correlated with robustness, except for 1st Neighbors (Figure 6D). This effect is particularly pronounced for DOMINO and ROBUST (bias-aware), both of which exhibit significantly reduced robustness in larger networks.

**Figure 6:**
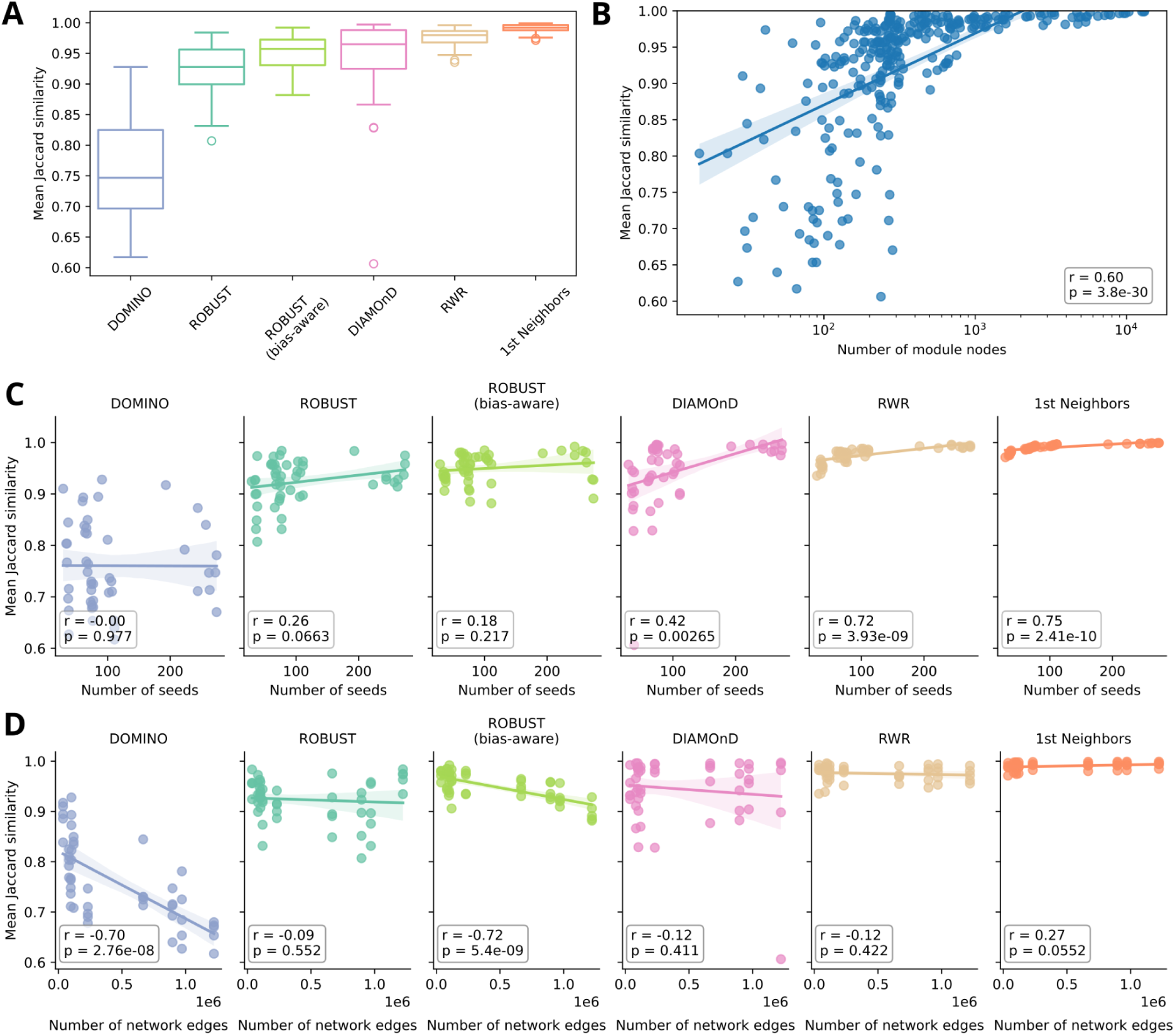
Robustness of AMIMs to leave-one-out perturbations of the input seed set. Robustness is quantified as the mean Jaccard similarity between the node set of the original module and those of its perturbed variants, with higher values indicating greater robustness. (**A**) Boxplots summarize results aggregated across different seed set–network combinations. (**B**–**D**) Correlations between robustness and module size, number of seed nodes, and input network size (measured by the number of edges). *r* indicates the Pearson correlation and *p* the corresponding p-value.

### Most AMIMs cannot reliably rediscover key disease genes

In addition to assessing robustness toward small input perturbations, the pipeline utilizes the leave-one-out seed perturbation to calculate a rediscovery rate, defined as the fraction of removed seeds that are reincluded in the module by the AMIM (see Methods) [30]. Since the seed genes are known to be associated with the corresponding disease, this metric provides an estimate of how effectively an AMIM might discover genes with previously unknown disease associations.

As shown in Figure 7A, the highest rediscovery rate is achieved by 1st Neighbors, which includes all direct interactors of the remaining seeds and reaches a median rediscovery rate above 50%. In contrast, the median rediscovery rates of other AMIMs are substantially lower, dropping below 10% for DOMINO, ROBUST, and ROBUST (bias-aware).

**Figure 7:**
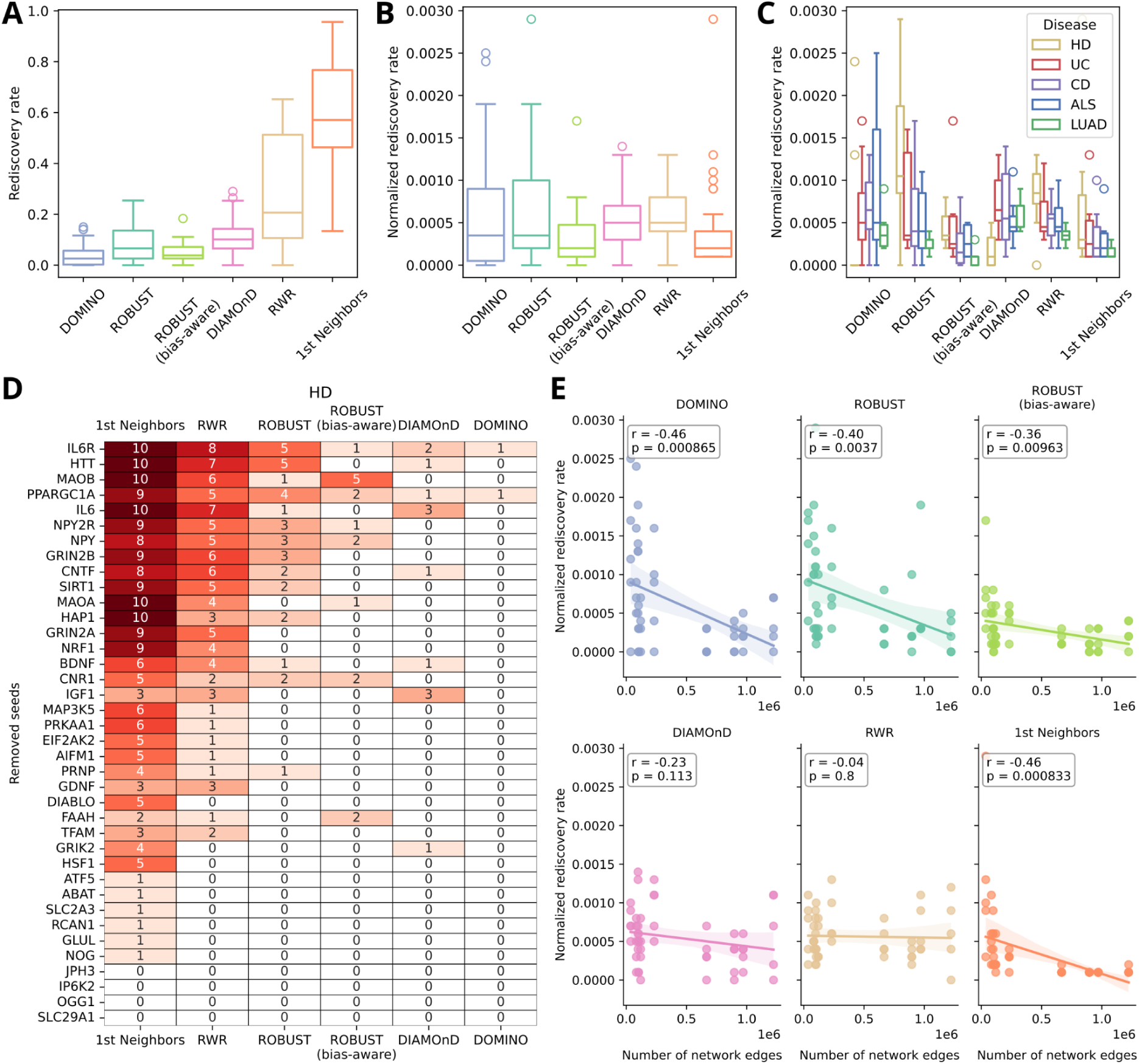
Results of the seed rediscovery analysis. (**A**) Distributions of seed rediscovery rates and (**B**) normalized seed rediscovery rates across different AMIMs. Boxplots summarize results aggregated across (**A**–**B**) different seed set–network combinations or (**C**) different networks. (**D**) Heatmap showing the number of times each AMIM successfully recovered each HD seed gene across different networks. As ten networks were used, this is the maximum possible value. Rows are sorted in decreasing order by row sum, and columns by column sum. (**E**) Correlation between normalized seed rediscovery rate and input network size (measured by the number of edges). corresponding p-value. *r* indicates the Pearson correlation and *p* the corresponding p-value.

However, 1st Neighbor modules tend to become very large (Figure 3A), and the probability of re-including a specific node increases simply by chance when module size grows. To account for this effect, the pipeline also reports a normalized rediscovery rate (Figure 7B), obtained by dividing the rediscovery rate by the number of nodes in the module (see Methods). After normalization, differences between methods are less pronounced, with DIAMOnD and RWR achieving the highest median normalized rediscovery rates, while ROBUST (bias-aware) and 1st Neighbors perform the worst. Notably, ROBUST (bias-aware) shows reduced performance relative to its original implementation, likely because disease-associated genes or proteins tend to be well- or even overstudied [41]. By penalizing such genes, ROBUST (bias-aware) reduces their inclusion, lowering rediscovery rates. This not only explains its reduced performance but also highlights a limitation of the rediscovery rate as a proxy for the ability to identify novel disease genes: methods may recover known disease genes via study-bias shortcuts that do not apply to the discovery of previously unknown associations.

We hypothesized that seed rediscovery might depend on the underlying disease, with some diseases being more easily recovered than others. However, this is not the case: no disease consistently shows the highest or lowest normalized rediscovery rates across methods (Figure 7C). For example, HD seeds are among the easiest to recover for some AMIMs but among the hardest for others. In contrast, network choice has a clear impact, with network size consistently showing a negative correlation with the normalized rediscovery rate (Figure 7E). This may again reflect study bias, as smaller, high-confidence networks tend to focus on well-studied proteins, which may make the rediscovery of well-characterized disease genes appear artificially easier.

Figure 7D illustrates the rediscovery of individual seed genes associated with HD, which the pipeline also reports. It enables the assessment of whether highly relevant genes are successfully recovered. HD is caused by pathogenic expansions of CAG trinucleotide repeats in the *HTT* gene [42]. We selected HD for closer examination due to its well-characterized genetic cause. Moreover, both the wild-type and mutant forms of *HTT* are known to interact with numerous proteins, resulting in a complex pathomechanism despite the disease’s monogenic origin [43]. Given its central role in HD pathogenesis, successful rediscovery of *HTT* by AMIMs would be expected. Indeed, *HTT* is identified as the second most frequently rediscovered HD seed gene. However, neither ROBUST (bias-aware) nor DOMINO recovered it at all, and ROBUST, RWR, and DIAMOnD only rediscovered it inconsistently across different network configurations.

Taken together, the seed rediscovery analysis demonstrates that while AMIMs have the potential to identify highly relevant genes, this is not guaranteed at the level of individual modules. Moreover, given the research bias toward well-studied disease genes, it is uncertain how well these findings translate to the discovery of previously unknown disease genes.

### Node inclusion becomes increasingly degree-driven for larger networks

Previous work by Lazareva et al. demonstrated that most AMIMs produce biologically meaningful results on real input networks that are similar to those on randomized networks, where the exact node degrees and thus hub nodes are preserved [16]. This finding suggests that many AMIMs primarily exploit node degrees, which are affected by study bias in the networks [41], rather than leveraging individual PPIs for module construction.

To detect this behavior within our pipeline, we implemented a similar analysis. Specifically, the input network is repeatedly rewired in a degree-preserving manner (see Methods), and the AMIMs are rerun on these randomized networks. The resulting modules are then compared to those derived from the original network using the same procedure as in the leave-one-out seed perturbation analysis. However, in this case, strong deviations from the original modules are desirable, as a lack of deviation would indicate that an AMIM primarily exploits node degree shortcuts.

Figure 8A summarizes the results of this analysis. Notably, AMIMs are far more strongly affected by network rewiring than by the removal of individual seed genes (median mean Jaccard similarity <0.3 compared to >0.9 for most AMIMs in the leave-one-out analysis, Figure 6A). This indicates that none of the evaluated AMIMs rely solely on node degrees. Among the methods, DOMINO is most strongly impacted by network randomization, consistent with the findings of Lazareva et al., who reported that DOMINO was the only AMIM to achieve significantly more biologically meaningful results on the original network than on the randomized ones. In contrast, the 1st Neighbors approach appears to rely most heavily on node degrees. This is expected, as high-degree nodes are more likely to interact with at least one seed node by chance. Furthermore, ROBUST shows a stronger dependence on node degrees compared to its bias-aware variant, providing evidence that ROBUST (bias-aware) successfully reduces the influence of research bias and degree-based shortcuts.

**Figure 8:**
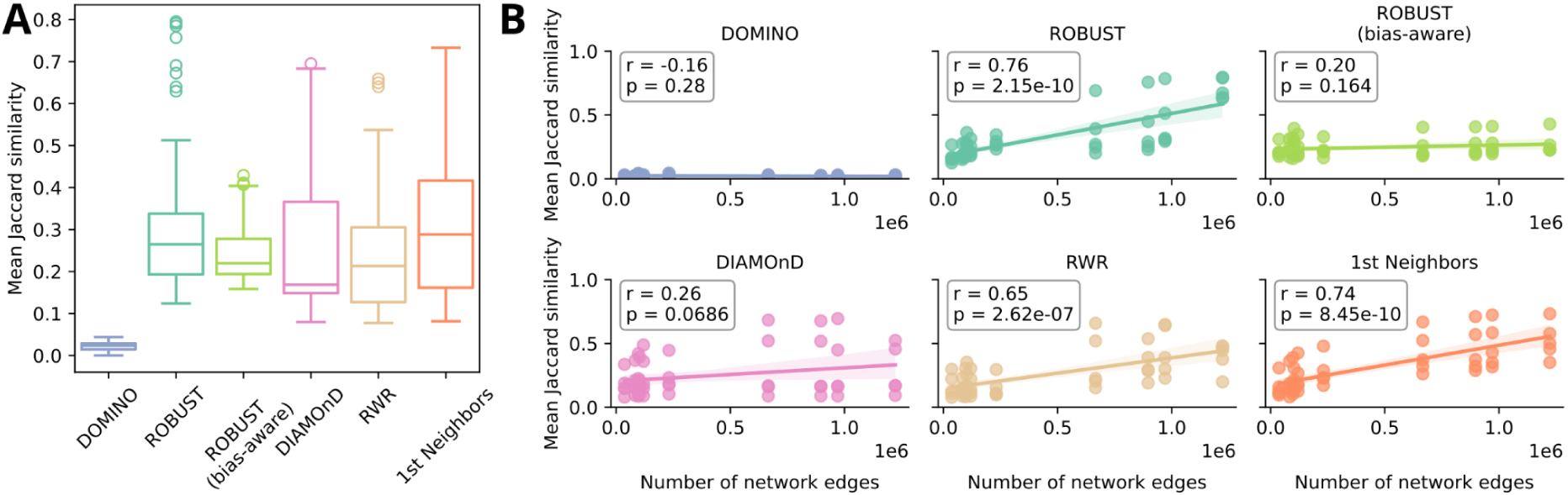
Results of the degree-perserving network rewiring analysis. High mean Jaccard similarities between the original modules and those based on the rewired networks, depicted on the y-axis, indicate a strong reliance on node degrees. (**A**) A single boxplot summarizes results aggregated across different seed set–network combinations. (**B**) Correlation between Jaccard similarity and input network size (measured by the number of edges). *r* indicates the Pearson correlation and *p* the corresponding p-value.

Interestingly, except for DOMINO, the AMIMs exhibit increased node-degree dependence with larger networks, which is statistically significant for ROBUST, RWR, and 1st Neighbors (see Figure 8B). This suggests that while the overall impact of node-degree bias on the AMIMs is only moderate, it becomes increasingly relevant for larger networks, and not all AMIMs are well-equipped to handle it effectively.

### Function-based module coherence depends on the individual case

The tool DIGEST was specifically developed to evaluate disease modules. It is based on the assumption that the genes within a module should be functionally coherent, i.e., involved in similar biological processes, represented through gene set annotations from Gene Ontology (GO) [44,45], including Biological Process (GO.BP), Cellular Component (GO.CC), and Molecular Function (GO.MF), as well as the Kyoto Encyclopedia of Genes and Genomes (KEGG) [46]. The pipeline integrates DIGEST in two different modes. The reference-free mode assesses the internal functional coherence between all module nodes, while the reference-based mode compares the functional coherence between the seed nodes and the added nodes.

In the reference-free mode (Figure 9A), the raw seed genes already exhibit strong functional coherence for GO.BP and KEGG, with coherence decreasing upon the inclusion of additional nodes. For GO.CC, only DIAMOnD and the 1st Neighbors approach improve upon the seed gene baseline. For GO.MF, significant results are observed only sporadically. In the reference-based mode (Figure 9B), the 1st Neighbors approach achieves the highest coherence across gene set sources. By contrast, ROBUST (bias-aware) yields modules with the lowest coherence in both modes, likely due to its inclusion of more distant genes (Figure 3D), which may be less functionally related. Overall, module coherence appears to depend strongly on the specific case, mode, and gene set source, without consistent patterns across different seed sets or input network configurations (Figure S5 and Figure S6).

**Figure 9:**
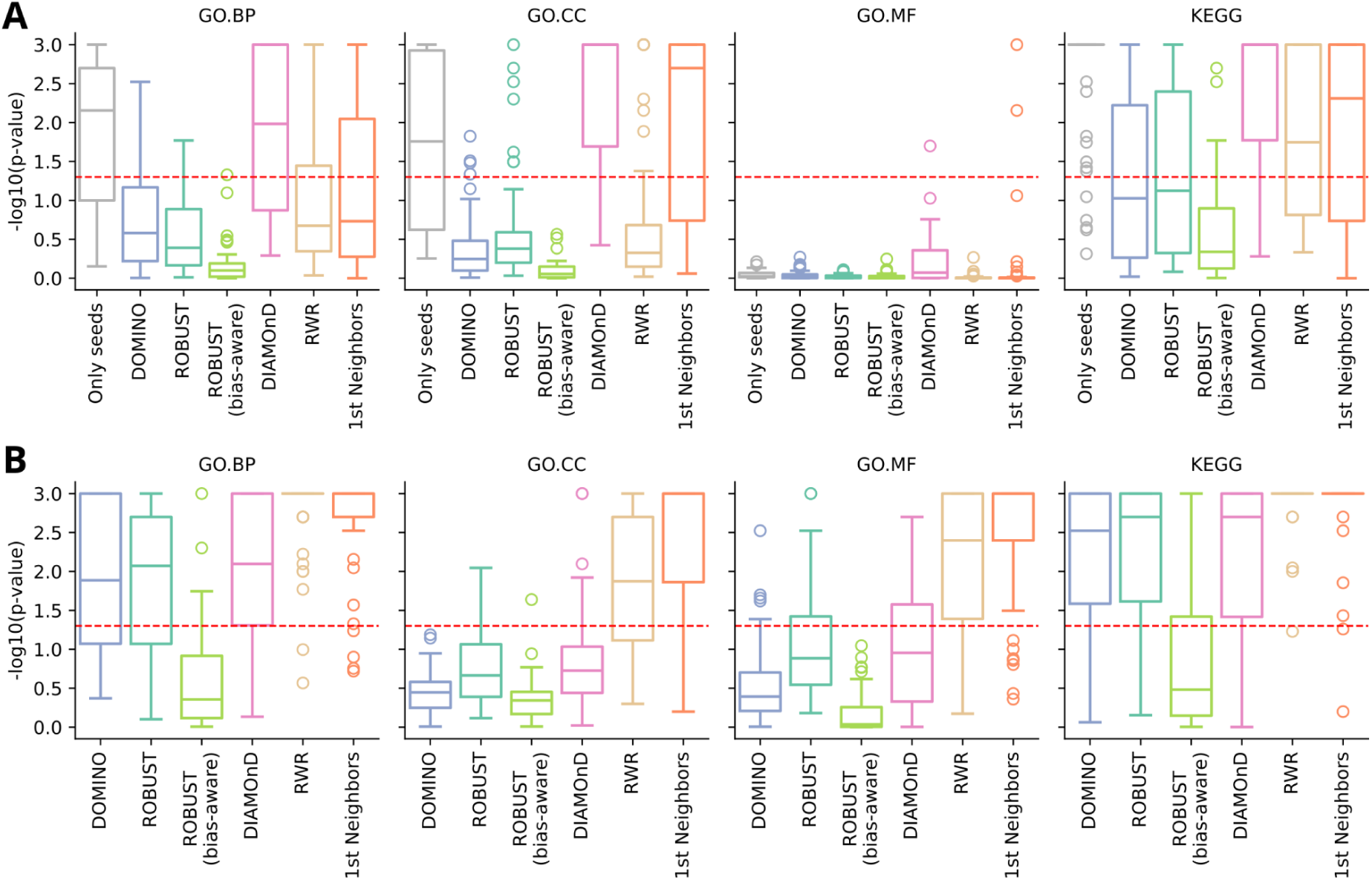
Results of the functional coherence analysis with DIGEST in (**A**) reference-free mode and (**B**) reference-based mode. Functional coherence is expressed through empirical p-values transformed using − *log*_10_. The significance level of 0.05 is indicated by the dashed line. A single boxplot summarizes the results for one AMIM in combination with one gene set source (GO.BP, GO.CC, GO.MF, or KEGG) aggregated across different seed set–network combinations.

## Discussion

### Method and network choice for module discovery

No single AMIM consistently outperformed the others in our demonstration. Instead, each showed distinct strengths and limitations that should be considered when selecting a method for a specific use case.

For example, in diseases involving multiple independent mechanisms, such as cancer [47], an AMIM capable of handling disconnected modules may be more appropriate than one that forces all seed nodes into a single component. Conversely, when working with a high-confidence set of seed nodes, it may be desired that the method retains all seeds in the final module. In contrast, for low-confidence seed sets that are likely to include false positives, a method that can filter out seeds that do not fit the context may be preferable. Similarly, if the seed nodes are expected to already capture the core mechanism, a method that only incorporates nearby nodes may be sufficient. In contrast, a more exploratory method could be advantageous when the initial seeds only partially represent the underlying biology.

Our analyses further underscored the critical influence of the input network choice. Although our findings do not support a specific network recommendation, we observed that larger networks are more frequently associated with undesirable behaviors when combined with certain AMIMs. For those, we anticipate smaller networks to yield more reliable results. Based on their analysis results, Yang et al. recommended the relatively small STRING network with a confidence threshold of 0.9, corresponding to the “highest confidence” configuration in our pipeline, over the larger BioGRID network, consistent with our own observations.

To guide users in selecting the most suitable approach, we summarize the outcomes of our analyses in Table 1, to enable an informed choice when combined with the evaluation results of their own runs.

**Table 1:** Summary of the AMIM properties revealed through the analysis performed to demonstrate the pipeline.

|  | **DOMINO** | **ROBUST** | **ROBUST**<br>**(bias-aware)** | **DIAMOnD** | **RWR** | **1st**<br>**Neighbors** |
| --- | --- | --- | --- | --- | --- | --- |
| Approach | Clustering | Connecting the seeds | Connecting the seeds | Expanding from the seeds | Expanding from the seeds and connecting the seeds | Expanding from the seeds |
| Designed for multiple, disconnected mechanisms? | Yes | No | No | Yes | No | Yes |
| Will retain all seed nodes in the module? (Figure 3C) | No | Yes, if part of the input network's LCC | Yes, if part of the input network's LCC | Yes | Yes, if part of the input network's LCC | Yes, if part of the input network's LCC |
| Adds nodes more distant from the seed nodes? (Figure 3D) | Yes, but very rarely | Yes, sometimes adds 2nd and 3rd neighbors | Yes, frequently | Yes, frequently adds 2nd and 3rd neighbors | Yes, sometimes adds 2nd and 3rd neighbors | No, only direct interactors |
| Are the modules specific to the seed set? (Figure 4) | Yes | Yes | Yes | Only for some seed sets | Only for some seed sets | Only for some seed sets |
| Are the modules specific to the seed set based only on the added nodes? (Figure 4) | Only for some seed sets | No | No | No | No | No |
| Are the modules impacted by network choice? (Figure 5) | Yes, heavily | Yes, heavily | Yes, heavily | Yes, moderately | Yes, moderately | Yes, moderately |
| Are the modules robust towards leave-one-out perturbations? (Figure 6) | No | Yes | Yes | Yes | Yes | Yes |
| Good at rediscovering left-out seeds? (Figure 7) | Average, if module size is considered | Average, if module size is considered | No | Yes, if module size is considered | Yes | Yes, but only if the module size is not considered |
| Bigger networks result in... | worse robustness; worse normalized | smaller modules; worse normalized | worse robustness; worse normalized |  | smaller modules; stronger dependence | bigger modules; worse normalized |
|  | seed<br>rediscovery | seed<br>rediscovery;<br>stronger<br>dependence<br>on node<br>degrees | seed<br>rediscovery |  | on node<br>degrees | seed<br>rediscovery;<br>stronger<br>dependence<br>on node<br>degrees |

### Effect of method-specific parameters on disease module topology

The topological properties of disease modules inferred by different AMIMs were obtained using default parameter settings (see Methods). Adjusting these parameters can directly affect several of the investigated properties. For DIAMOnD, module size is determined by the number of added nodes (*n*), and the seed weight (α) will influence explorativeness. In DOMINO, module size depends on the confidence thresholds (`slices_threshold` and `module_threshold`). For ROBUST, size is affected by the node inclusion prize (α) and the tree fraction threshold (θ). In its bias-aware variant, the study-bias penalty (γ) can be reduced to yield modules closer to the original implementation. For RWR, a trade-off between local and global exploration of the seed’s neighborhood is determined by the restart probability *r*.

### Limitations and Outlook

We found that the AMIMs in our pipeline produced comparable modules when a seed gene is left out, but struggled to consistently recover the left-out seed and to produce disease-specific modules independent of the input network choice. This suggests that the AMIMs cannot always reliably distinguish between truly disease-relevant nodes and background noise in the underlying networks. In the future, we plan to leverage our modular pipeline design to integrate additional AMIMs [38,48] and evaluate whether alternative algorithms can improve performance. However, the lack of performance in these categories could also stem from both excessive noise or incompleteness in the input networks or seed sets [49,50], which may not be resolvable through algorithmic improvements alone. For both input types, there is an inherent trade-off between increased false-positive rates in larger network and seed set sizes and increased false-negative rates in smaller ones. All AMIMs currently implemented in the pipeline use binary seed weights and network edges, which we focused on due to their flexibility, simplicity, and performance in previous studies [11]. A possible way to address this trade-off between input size and accuracy in the future would be to integrate methods that use continuous node weights [51,52], or weighted network edges [34,36], for example, by using edge-dependent transition probabilities in an RWR approach, enabling direct representation of confidence levels in disease associations or protein interactions. We also aim to include approaches that use multi-omics datasets [53] or directed network edges [54]. Furthermore, the input networks provided through our pipeline interface are context-agnostic and do not account for tissue-specific variations in PPI interactions. While users can already provide custom, context-specific networks, we plan to add functionality that automatically filters PPIs based on genes or proteins expressed in a specified tissue of interest, leveraging expression databases [55,56].

While the evaluation procedures in our pipeline provide insights into the reliability of the inferred disease modules, these should be considered only as indicative evidence, since a complete ground truth would be required to assess AMIM performance definitively. Moreover, certain evaluation steps may be subject to research bias and circular reasoning. For example, seed sets often consist of well-studied disease genes or proteins with many known PPIs, making them easier to rediscover using network-based approaches. Additionally, these genes or proteins may have already been analyzed together in the context of specific diseases or pathways, potentially biasing the assessment of functional coherence or pathway enrichment analyses. The underperformance of the research-bias-aware implementation of ROBUST in these analyses further supports the notion that evaluation procedures relying on prior knowledge are susceptible to such biases.

One of the advantages of reusable pipelines that utilize containers for software deployment is reproducibility. In our pipeline, full reproducibility is partially limited by randomization-based evaluation procedures, such as DIGEST or network rewiring, as well as by AMIMs like ROBUST or DOMINO, which rely on heuristics and are not fully deterministic. Additionally, some of the queried resources, such as the NeDRex database and Drugst.One, are continuously updated, which will result in changes over time. While the rewired networks generated by the pipeline can be reused in subsequent analyses to ensure reproducibility at this step, overall, two runs of the pipeline on identical input are not guaranteed to yield entirely identical results.

Additional limitations of our pipeline include the current inability to explore different tool parameter configurations or tunings within a single run. Furthermore, input perturbation-based evaluation procedures can be computationally intensive and may only be feasible with sufficient resources, such as a compute cluster. Finally, algorithmically inferred disease modules often require curation by biomedical experts, which cannot be fully automated. To facilitate this process, we provide Drugst.One export links, which, together with its recent extension, Drugst.One DREAM [57], supports module refinement by offering additional context and an intuitive web interface.

In addition to the previously mentioned extensions concerning the included AMIMs, supported input types, and tissue-specific networks, we plan to implement dedicated evaluation procedures for the generated drug rankings [31], enable the selection of different seed sets from popular databases directly through the pipeline interface (similar to the network selection), expand the leave-one-out analysis with an option to remove multiple seeds at a time, and generate a pre-computed atlas of disease modules covering a wide range of diseases.

## Conclusions

The presented pipeline provides access to a diverse collection of disease module discovery algorithms within the most comprehensive evaluation framework to date. Through efficient parallelization, it enables perturbation-based evaluation procedures and scales seamlessly to large input datasets. Applied to 50 input combinations, the pipeline revealed characteristic strengths of the included methods while also highlighting limitations, such as strong dependence on the input network and only moderate performance in seed rediscovery analyses. Built on a modular and extensible code base, it is designed to serve as a sustainable platform for evaluating new methods and networks, thereby supporting systematic benchmarking and tracking progress in the field.

## Materials and methods

### Pipeline

#### Implementation

The pipeline is implemented using Nextflow [23] with its DSL2 extension, which supports a modular design for improved maintainability, configurability, and future expandability. Nextflow manages the execution of all pipeline steps and parallelizes processes where possible. It caches the results of individual steps, enabling resumption from the last successful point in case of failure. This also allows for running specific parts of the pipeline and incorporating additional steps later without needing to recompute previous results. Software dependencies are automatically deployed through Docker or Singularity [58], requiring the user to install only Nextflow and a compatible container runtime. Nextflow pipelines are highly portable, allowing them to run on HPC clusters as well as various cloud computing platforms.

Our pipeline is part of the nf-core project [24] and uses the nf-core template, adhering to best practices for code structure and documentation. Continuous integration (CI) tests monitor the pipeline development, and an included test dataset enables users to easily verify a successful installation.

Custom analysis scripts are written in Python (v3.12) and are centered around the *graph-tool* library [59] (https://graph-tool.skewed.de/, v2.77) for network analysis and its binary GT file format and for efficiently passing graph data between processes. These dependencies are deployed via a shared container (https://github.com/REPO4EU/modulediscovery_python_dependencies). Most external tools are run in dedicated containers for better modularity and to prevent software version conflicts.

#### Input

The main inputs to the pipeline are a text file containing the seed nodes (one per line) and a network file, which can be in CSV, GT, GraphML, or DOT format. It is also possible to provide multiple files for each input type. In this case, the pipeline can either run on all possible input combinations or on specific pairs defined in a sample sheet. Seed files are filtered to include only nodes present in the corresponding network file. If multiple networks are provided, the pipeline generates a separately filtered seed file for each network.

Some tools further require specifying the ID space of the input. Supported ID spaces are HGNC Symbols [60], Entrez IDs [61], or Ensembl IDs [62] for genes and UniProt accession numbers (AC) [63] for proteins.

#### Available networks

Instead of supplying their own input network, users can select from a variety of widely used human PPI networks, including STRING [34], BioGRID [35], HIPPIE [36], IID [37], and NeDRex [31]. For several of the sources, we provide multiple subsets, resulting in a total of ten network options from five databases. For STRING, users can choose between networks that include or exclude non-physical functional interactions, each available with two confidence thresholds (score > 0.9 or > 0.7). For HIPPIE, both medium- and high-confidence versions are available, based on the score cutoffs of 0.63 and 0.73, respectively, as defined on the HIPPIE website (https://cbdm-01.zdv.uni-mainz.de/∼mschaefer/hippie/information.php). For NeDRex, we only included experimentally validated interactions between reviewed proteins. We provide a standard and a high-confidence version, the latter filtered by a method score greater than 13.5 [64].

The networks were downloaded from their respective primary sources, using UniProt accession numbers (ACs) as ID space whenever available. HIPPIE and STRING did not provide UniProt ACs directly; instead, they used UniProt entry names and Ensembl protein IDs, respectively. These identifiers were converted to UniProt ACs using UniProt’s official ID mapping file (https://ftp.uniprot.org/pub/databases/uniprot/current_release/knowledgebase/idmapping/by_organism/HUMAN_9606_idmapping_selected.tab.gz, accessed on 18.03.2025).

The mapping file provides a one-to-one correspondence between UniProt entry names and UniProt ACs, but may result in a many-to-many mapping between Ensembl protein IDs. When an original identifier mapped to multiple UniProt ACs, we created a separate network node for each resulting UniProt AC and assumed each to interact with all partners of the original identifier. Conversely, when multiple original identifiers mapped to the same UniProt AC, the corresponding nodes were merged.

The resulting UniProt AC-based networks were subsequently mapped to the other supported ID spaces. Ensembl and Entrez gene IDs were obtained using the same UniProt mapping file, while HGNC symbols were retrieved using the *mygene* Python package (https://github.com/biothings/mygene.py, v3.2.2), as they were not included in the UniProt mapping file. Multi-mapper IDs were handled as outlined above. Duplicate edges and self-loops were removed from all networks.

The code for downloading, parsing, and mapping all networks is available at https://github.com/REPO4EU/network_preparation. The process is fully automated, and both download sources and parameters can be configured via a settings file to easily add or modify data sources for future releases.

To use the available networks, instead of a file path, the user can specify a keyword to automatically load the network (see Table S1).

#### Integrated AMIMs

The pipeline currently incorporates six disease module discovery methods, chosen based on their popularity, code availability, and ease of integration. Default parameter values are derived from each method’s recommended or standard settings and can either be configured directly through pipeline parameters or modified by supplying custom command-line options to the respective tools. The included methods are:

**DOMINO** (Discovery of Modules In Networks using Omic) [11] begins by partitioning the network into disjoint slices using Louvain clustering [65], selecting those enriched for seed nodes based on a hypergeometric test. The selected slices are refined by solving the Prize Collecting Steiner Tree (PCST) problem [13] and are then further subdivided into putative modules, each containing no more than ten nodes. These modules are again tested for seed enrichment using a hypergeometric test. DOMINO outputs a flexible number of modules, which may belong to the same or to different connected components. In the pipeline, all returned modules are merged into a single module by taking their union. Each node is annotated with an additional identifier indicating its original DOMINO module. DOMINO (https://github.com/Shamir-Lab/DOMINO, v1.0.0) is deployed in the pipeline via its available biocontainer (https://quay.io/repository/biocontainers/domino). The method provides parameters to adjust the significance thresholds for slices (`slices_threshold`) and putative modules (`module_threshold`), which default to 0.3 and 0.05, respectively.

The **DIAMOnD** (DIseAse MOdule Detection) algorithm [2] iteratively expands an initial set of seed nodes by adding one node at a time. At each step, it selects the node with the highest connectivity significance to the current seed set, computed using a hypergeometric test. This process continues until a predefined number of nodes have been added. DIAMOnD only returns a list of added nodes, so the pipeline combines these with the seed nodes to produce the final output module. The result is a single disease module, which may consist of one or multiple connected components. DIAMOnD (https://github.com/dinaghiassian/DIAMOnD) is deployed in the pipeline via a custom container (https://hub.docker.com/r/djskelton/diamond) originally created for the NeDRex platform. The total number of added nodes *n* and the weight assigned to the initial seeds α are configurable via parameters and default to 200 and 1.0, respectively, in accordance with the authors’ recommendations.

**ROBUST** (robust disease module mining via enumeration of diverse prize-collecting Steiner trees) [12] repeatedly connects seed genes by solving the PCST problem. In each iteration, nodes included in previous solutions are penalized, reducing their likelihood of being selected again. The final disease module comprises nodes that have appeared in a sufficient number of these solutions to increase robustness. It may consist of one or multiple connected components. ROBUST (https://github.com/bionetslab/robust) is deployed in the pipeline via a custom container (https://hub.docker.com/r/djskelton/robust) originally created for the NeDRex platform. The initial value of integrating non-seed nodes α, value reduction factor β, number of PCST *n*, and the fraction of PCST runs a node must appear in to be included in the final module τ can all be configured via parameters. By default, these are set to (α, β, *n*, τ) = (0. 25, 0. 9, 30, 0. 1), based on the authors’ recommendations.

**ROBUST (bias-aware)** [14] adopts the same strategy as ROBUST but increases the edge costs for nodes frequently used as baits in PPI detection experiments. This penalization is designed to counteract study bias in current PPI networks [41]. ROBUST (bias-aware) (https://github.com/bionetslab/robust_bias_aware, v.0.0.1) is deployed in the pipeline via its available biocontainer (https://quay.io/repository/biocontainers/robust-bias-aware). The method provides the same parameters as ROBUST, along with an additional γ parameter (default: 1.0), which can be used to adjust the study-bias edge penalty.

**RWR** (Random Walk with Restart) [15] models signal diffusion on a network by simulating a walker that moves randomly between connected nodes but, with a restart probability *r*, returns to the seed nodes. This parameter balances global exploration of the interactome with local focus around the seeds, and the resulting steady-state probabilities provide a ranking of proteins by their relevance to the seeds. The size of the disease module is determined by including nodes ranked by the RWR until all seeds are connected, which will result in a module consisting of exactly one connected component. RWR is deterministically implemented (no simulation). We set *r* to 0.8 to have an effective trade-off between local and global exploration of the seeds’ neighborhood [66]. The method can scale the nodes’ visiting probability by the square root of their degree using the `scaling` parameter (default: False) and allows the use of the symmetrical Markov matrix with the `symmetrical` parameter (default: False).

**1st Neighbors** [9] includes every network node that directly interacts with at least one seed, resulting in a module that may contain one or more connected components. The pipeline implements this approach using the *graph-tool* library.

Additionally, the pipeline integrates a pseudo-AMIM that outputs only the seed nodes along with the edges connecting them within the input network. Filtering the seed genes that are not present in the input network, this configuration serves as a baseline for comparison with other AMIMs and is referred to as “**Only seeds**” throughout this manuscript.

#### Module topology

Let *N* = (*V*, *E*) be the input network with node set *V* and edge set *E*, *S* the seed node set with *S* ⊆ *V*, and *M*(*S*, *N*) ⊆ *V* the set of nodes in a disease module inferred with seeds *S* and network *N*. All topological measures for a disease module are calculated on its induced subnetwork *N*[*M*(*S*, *N*)] with node set *M*(*S*, *N*) and edge set consisting of all edges in *E* which connect two nodes included in *M*(*S*, *N*).

The pipeline reports topological properties for each module, including the number of nodes and edges, the number of included seeds, the maximum distance from any added node to its nearest seed, the pseudo diameter (a heuristic estimate of the longest shortest path between any two nodes), the number of connected components, the size of the largest connected component, and the number of isolated nodes, i.e., nodes without any connections to others. The maximum distance from any added node *a* to its nearest seed *s* is calculated as *max*_*a*∈*M*(*S*,*N*)\*S*, *s*∈*S*_ *dist*(*a*, *s*), where *dist*(*a*, *s*) is the length of the shortest path between nodes *a* and *s* in the subnetwork induced by the node set *M*(*S*, *N*). The pseudo diameter is calculated using the *pseudo_diameter* function in the *graph-tool* library.

#### Module overlap

To evaluate similarities between modules produced through different seed sets, networks, or AMIMs, the pipeline calculates pairwise overlaps between all modules. This includes the number of shared nodes |*A* ∩ *B*| between two module node sets *A* and *B*, and their Jaccard similarity, i.e. 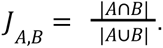 To focus only on added nodes, the same measures are also computed on *A*\*S* and *B*\*S*, where *S* is the set of seed nodes.

#### Biological module evaluation

Over-representation analysis is performed using g:Profiler through its R package *gprofiler2* (https://cran.r-project.org/web/packages/gprofiler2/index.html, v0.2.2) [29], which is integrated into the pipeline via the corresponding nf-core module (https://nf-co.re/modules/gprofiler2_gost/). The pipeline performs over-representation analysis using module nodes as the foreground and all network nodes as the background. By default, it considers gene sets from Gene Ontology (GO) [44,45], WikiPathways [67], Reactome [68], and KEGG [46].

Functional coherence of the modules is assessed using DIGEST [17] (https://github.com/bionetslab/digest, v0.2.16), which the pipeline integrates via its biocontainer (https://quay.io/repository/biocontainers/biodigest). The pipeline runs DIGEST in two modes: a reference-free mode (`mode=“subnetwork”`), which evaluates the functional coherence of all nodes within a module, and a reference-based (`mode=“subnetwork-set”`) mode, which assesses the coherence between the seed nodes and the nodes added during module construction. Both modes use Jaccard similarity as a distance metric and rely on the pipeline input network(s) to generate 1,000 random modules for perturbation-based significance testing.

#### Input perturbation-based evaluation

##### Seed perturbation

To evaluate the robustness of module discovery methods against minor perturbations of the seed set, the pipeline performs a leave-one-out analysis. Given an initial seed set 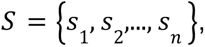 it generates perturbed sets 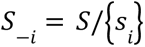 by systematically removing each seed node *s_i_*. For each perturbed set *S*_-*i*_ and corresponding network *N*, a module with node set *M*(*S*_-*i*_, *N*) is computed and compared to the original node set of the original module *M*(*S*, *N*).

The robustness for each perturbation run is measured through the Jaccard similarity, i.e., 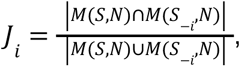 and summarized across runs using the mean.

The rediscovery rate [30] is calculated as the fraction of omitted seeds that are reincluded in the corresponding run, i.e., 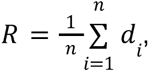 where *d* is 1 if *s* ∈ *M*(*S*_-*i*_, *N*) and 0 otherwise. Since larger modules will have a higher chance of randomly reincluding an omitted seed, the rediscovery rate is normalized to account for the size of the original module, i.e., 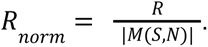

##### Network perturbation

The reliance of module discovery methods on node degree is evaluated by randomly rewiring the network while preserving the original degree of every node. For a network *N* defined through a set of nodes *V* and edges *E*, this is accomplished by iteratively selecting four nodes *a*, *b*, *c*, *d* ∈ *V* with (*a*, *b*) ∈ *E* and (*c*, *d*) ∈ *E*, and replacing the edges with (*a*, *d*) and (*b*, *c*), provided that this operation does not introduce self-loops or duplicate edges [69]. The network rewiring is performed using the *graph-tool* function *random_rewire* with `“constrained-configuration”` as `model` and 100 full sweeps over all edges. The number of perturbed networks *m* can be specified by the user and defaults to 100. Modules with node sets *M*(*S*, *N_j_*) are inferred for every perturbed network *N_j_*. The perturbation impact is again measured using Jaccard similarity, i.e., 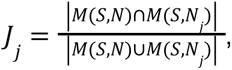, and summarized across all perturbed networks using the mean.

#### Annotation and BioPAX format

To characterize the inferred disease modules functionally, we perform an automated annotation step using information from NeDRexDB [31], a knowledge graph integrating a range of primary databases for network medicine-based drug repurposing. The annotated modules are represented through the standardized BioPAX [32] Level 3 format. The publicly available NeDRex API is queried based on the genes or proteins included in the disease module. If the module nodes are represented by gene-level identifiers, the corresponding protein products are annotated. Conversely, if protein-level identifiers are used, the associated coding genes are annotated. Disorders are linked to genes according to known associations. For proteins targeted by drugs, the corresponding drugs are retrieved together with their known side effects. Based on the obtained drugs and disorders, indications and contraindications of drugs for disorders are incorporated. Moreover, cellular components associated with the proteins are integrated based on GO annotations [44,45]. The *pybiopax* Python package (https://github.com/gyorilab/pybiopax, v0.1.4) [70] is used to parse and process BioPAX data. Since BioPAX Level 3 does not natively support entities for disorders and side effects, these are included as external references.

#### Drugst.One export and drug prioritization

For each inferred disease module, the pipeline generates a hyperlink enabling interactive exploration via the network medicine web tool Drugst.One [18]. These links are included in the pipeline report. Additionally, the associated Python package, *drugstone* (https://github.com/drugst-one/python-package, v0.4.5), is used to identify potential drug candidates that target module nodes. To prioritize compounds, the pipeline integrates three network-based ranking algorithms (see Figure 1B): degree centrality, which ranks compounds by the number of their targets within the module; harmonic centrality, which considers the average shortest distance from each compound to all module nodes; and TrustRank, ranking compounds based on network propagation [71]. Further details on these methods can be found in the supplementary material of the Drugst.One publication [18].

#### Network visualizations and pipeline report

Visual network representations of the inferred modules (both with and without assigned drugs) are generated using the graph-tool package and provided in PNG, SVG, and PDF formats. Additionally, interactive HTML visualizations are created with the pyvis package (https://github.com/WestHealth/pyvis, v0.3).

The pipeline results are summarized through an HTML report created with MultiQC (https://github.com/MultiQC/MultiQC, v1.27) [33]. MultiQC is a standard component of many nf-core pipelines, providing overviews of tool outputs, software versions, and execution commands. As MultiQC does not natively support the tools in our pipeline, they are integrated using MultiQC’s functionality to incorporate custom content.

### Pipeline demonstration

To demonstrate the pipeline, disease-associated genes from five complex diseases were used as seed sets. These genes were obtained from DisGeNET (https://www.disgenet.com/, v24.3) [25]. Table S2 lists the specific disease terms along with the number of associated genes included for each. The demonstration utilized all ten networks integrated into the pipeline (see Table S1) and was executed in a single run (see Supplementary Note 1).

The pipeline was executed using Nextflow (v24.10.05) on a high-performance computing (HPC) cluster running Ubuntu (v22.04.5), using approximately 2,300 CPU hours. Software dependencies were managed using Singularity (v3.8.7) [58].

Downstream analyses and visualizations were performed in Python (v3.12) using the *seaborn* package (https://github.com/mwaskom/seaborn, v0.13) [72]. Correlations between continuous variables were computed using Pearson’s correlation, as implemented in `scipy.stats.pearsonr` (https://github.com/scipy/scipy, v1.14) [73]. Linear regression lines were fitted and plotted using *seaborn’s lmplot* function. Hierarchically clustered heatmaps were generated with *seaborn’s clustermap* function using the `“ward”` linkage method. The order of columns and rows was randomly shuffled prior to clustering to eliminate potential biases arising from the initial ordering.

## Author contributions

**J.K.** (Conceptualization [equal], Data curation [lead], Formal analysis [lead], Investigation [lead], Methodology [lead], Software [lead], Validation [lead], Visualization [lead], Writing – Original Draft [lead], Writing – Review & Editing [equal]), **C.B.** (Conceptualization [support], Methodology [support], Software [support], Validation [support], Writing – Original Draft [support], Writing – Review & Editing [equal]), **L.M.S.** (Software [support], Validation [support], Writing – Review & Editing [equal]), **J.A.** (Conceptualization [support], Methodology [support], Software [support], Validation [support], Writing – Review & Editing [equal]), **Q.M.** (Conceptualization [support], Methodology [support], Software [support], Writing – Review & Editing [equal]), **T.P.** (Software [support], Writing – Review & Editing [support]), **M.T.** (Software [support], Writing – Review & Editing [support]), **F.M.D.** (Conceptualization [support], Validation [support], Writing – Review & Editing [support]), **C.N.** (Conceptualization [support], Writing – Review & Editing [support]), **H.H.H.W.S.** (Conceptualization [support], Project administration [support], Writing – Review & Editing [support]), **J.M.** (Conceptualization [support], Funding acquisition [support], Supervision [support], Writing – Review & Editing [support]), **A.M.** (Conceptualization [support], Methodology [support], Writing – Review & Editing [equal]), **J.B.** (Funding acquisition [support], Supervision [support], Writing – Review & Editing [equal]), **E.G.** (Conceptualization [support], Funding acquisition [support], Project administration [support], Supervision [support], Writing – Review & Editing [equal]), **M.L.** (Conceptualization [equal], Methodology [support], Funding acquisition [lead], Project administration [lead], Resources [lead], Supervision [lead], Writing – Review & Editing [equal]).

## Acknowledgements

Funded by the European Union. Views and opinions expressed are however those of the author(s) only and do not necessarily reflect those of the European Union or the European Research Executive Agency. Neither the European Union nor the granting authority can be held responsible for them. This work was also partly supported by the Swiss State Secretariat for Education, Research, and Innovation (SERI) under contract No. 22.00115. Funded by the Deutsche Forschungsgemeinschaft (DFG, German Research Foundation) [422216132]. We thank Lars J. Jensen for his insightful discussions and for suggesting the combination of weighted network edges with a random walk approach. ChatGPT and Grammarly were used to improve the readability and language of the manuscript. All generated text was reviewed and edited by the authors, who take full responsibility for the final content.

## Data availability

- Pipeline code: GitHub (https://github.com/nf-core/diseasemodulediscovery)
- Analysis scripts for the pipeline demonstration: GitHub (https://github.com/REPO4EU/modulediscovery_demonstration)
- Input network preparation code: GitHub (https://github.com/REPO4EU/network_preparation)
- Prepared input networks: Zenodo (https://zenodo.org/records/17258762)
- Results of the pipeline demonstration run: Zenodo (https://zenodo.org/records/17536307)

