## Supplement for "Inferring and Evaluating Network Medicine-Based Disease Modules with Nextflow"

| Keyword | Version | Nodes | Edges | Description |
|---|---|---|---|---|
| `string_min900` | v12.0 | 11,971 | 93,559 | Human PPI network obtained from STRING, including physical and functional interactions with a score greater than 0.9. |
| `string_min700` | v12.0 | 15,788 | 224,045 | Human PPI network obtained from STRING, including physical and functional interactions with a score greater than 0.7. |
| `string_physical_min900` | v12.0 | 7,722 | 34,141 | Human PPI network obtained from STRING, including physical interactions with a score greater than 0.9. |
| `string_physical_min700` | v12.0 | 10,465 | 78,878 | Human PPI network obtained from STRING, including physical interactions with a score greater than 0.7. |
| `biogrid` | 4.4.242 | 18,101 | 865,553 | Human PPI network obtained from BioGRID. |
| `hippie_high_confidence` | v2.3 | 13,246 | 112,202 | Human PPI network obtained from HIPPIE, including only high-confidence interactions with a score greater than 0.73. |
| `hippie_medium_confidence` | v2.3 | 16,613 | 637,499 | Human PPI network obtained from HIPPIE, including only interactions with a score greater than 0.63. |
| `iid` | 2025-03-18 | 19,598 | 1,202,716 | Human PPI network obtained from IID. |
| `nedrex` | 2025-03-18 | 18,718 | 935,139 | Human PPI network queried from NeDRexDB, including only experimentally validated interactions. |
| `nedrex_high_confidence` | 2025-03-18 | 12,827 | 95,944 | Human PPI network queried from NeDRexDB, including only experimentally validated interactions with a score greater than 13.5. |

***Table S1:*** *Overview of PPI networks accessible via the pipeline interface. For sources without versioning, the query date is provided. Node and edge counts are based on UniProt AC IDs.*

| Abbreviation | Full name | DisGeNET term | Number of genes |
|---|---|---|---|
| HD | Huntington’s disease | Huntington Disease, C0020179 | 40 |
| UC | Ulcerative colitis | Ulcerative Colitis, C0009324 | 76 |
| CD | Crohn’s disease | Crohn's disease of large bowel, C0156147 | 78 |
| ALS | Amyotrophic lateral sclerosis | Amyotrophic Lateral Sclerosis, C0002736 | 127 |
| LUAD | Lung adenocarcinoma | Adenocarcinoma of lung, C0152013 | 280 |

***Table S2:*** *Overview of the seed gene sets used for the pipeline demonstration*

**Supplementary Note 1**

The command that was used to run the pipeline demonstration:

```
nextflow run nf-core/diseasemodulediscovery \
-r 10d49e1a3808f1046c181d1e6b7dac1481f2bdb5 \
-profile daisybio,singularity,keep_work \
--id_space symbol \
--seeds
../../data/seeds/ALS.tsv,../../data/seeds/CD.tsv,../../data/seeds/HD.tsv
,../../data/seeds/LUAD.tsv,../../data/seeds/UC.tsv \
--network
string_min900,string_min700,string_physical_min900,string_physical_min70
0,biogrid,hippie_high_confidence,hippie_medium_confidence,iid,nedrex,ned
rex_high_confidence \
--run_seed_permutation \
--run_network_permutation \
--outdir results
```

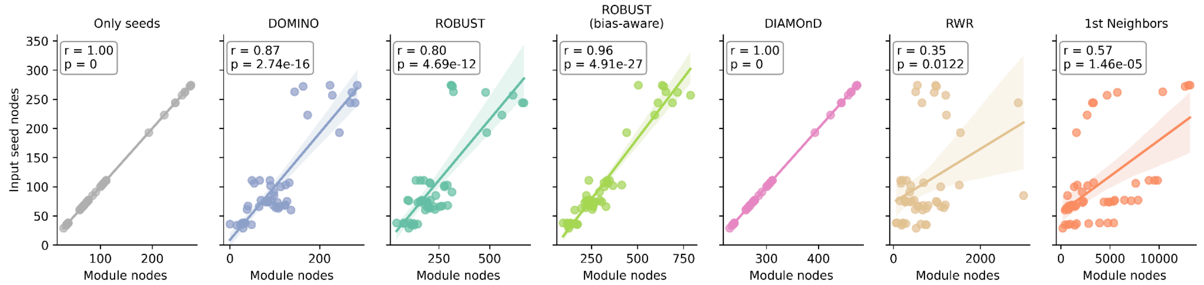


***Figure S1:** Correlation between the number of module nodes and the number of seed nodes used. $r$ indicates the Pearson correlation and $p$ the corresponding p-value.*

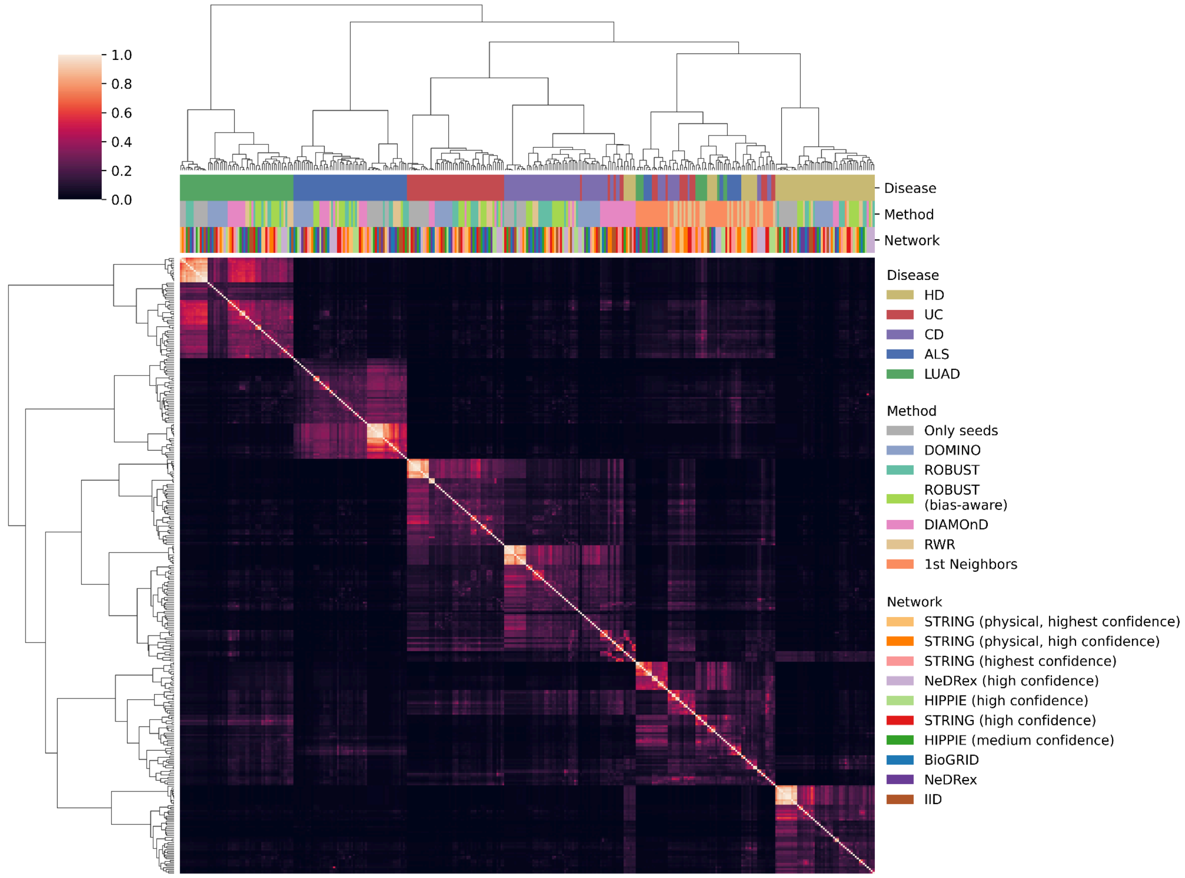


***Figure S2:** Heatmap and hierarchical clusterings based on the pair-wise node set similarities (measured through the Jaccard similarity) of modules inferred using different diseases, networks, and methods. All module nodes are considered.*

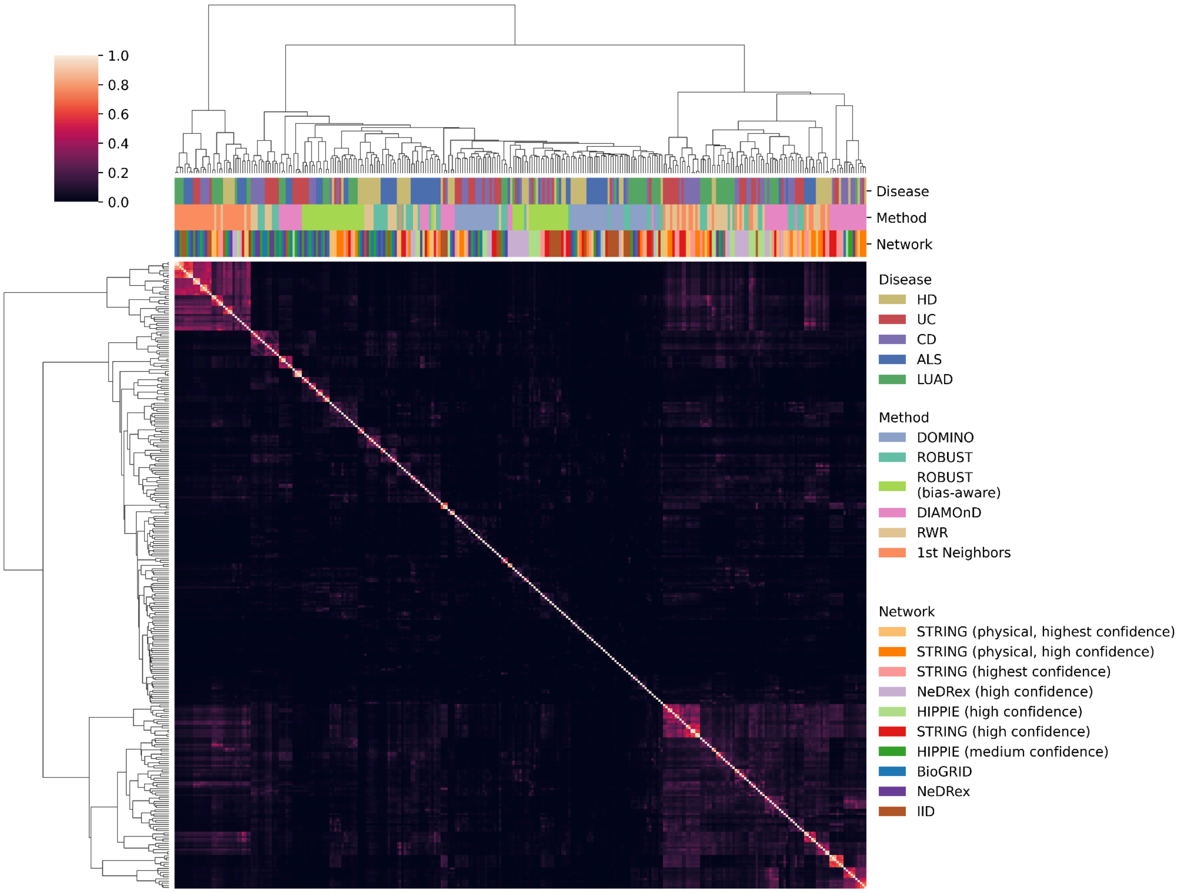


***Figure S3:*** *Heatmap and hierarchical clusterings based on the pair-wise node set similarities (measured through the Jaccard index) of modules inferred using different diseases, networks, and methods. Only added nodes (no seed nodes) are considered for the overlap calculation.*

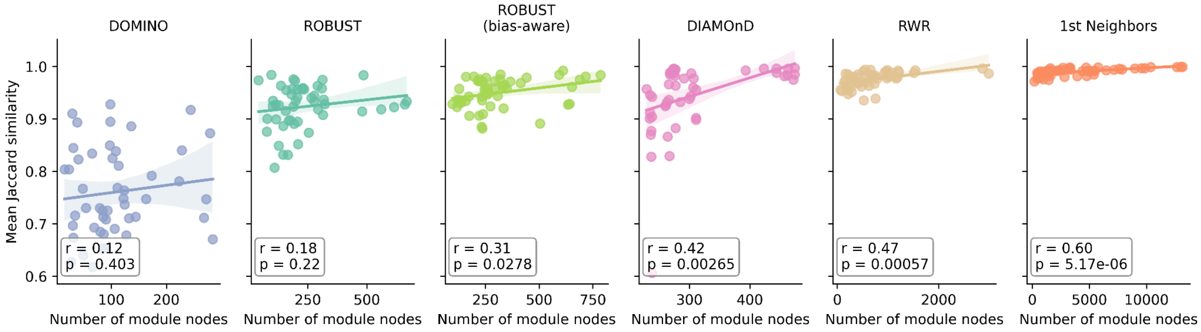


***Figure S4:*** *Correlations between robustness and module size (expressed through the number of included nodes) for different AMIMs.* $r$ *indicates the Pearson correlation and* $p$ *the corresponding p-value.*

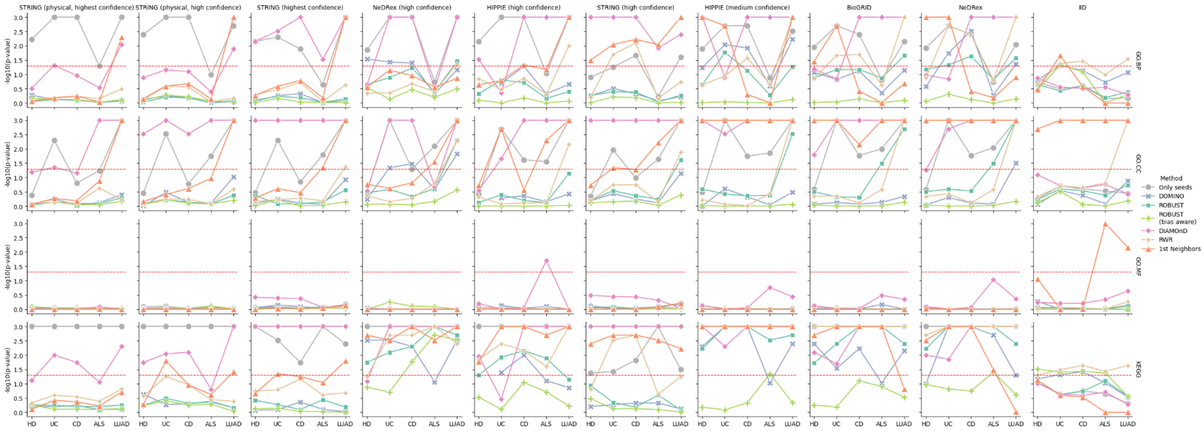


***Figure S5:*** *Results of the functional coherence analysis with DIGEST in reference-free mode for different AMIM, network, gene set source (GO.BP, GO.CC, GO.MF, or KEGG), and disease combinations. Functional coherence is expressed through empirical p-values transformed using* $-\log_{10}$. *The significance level of 0.05 is indicated by the dashed line.*

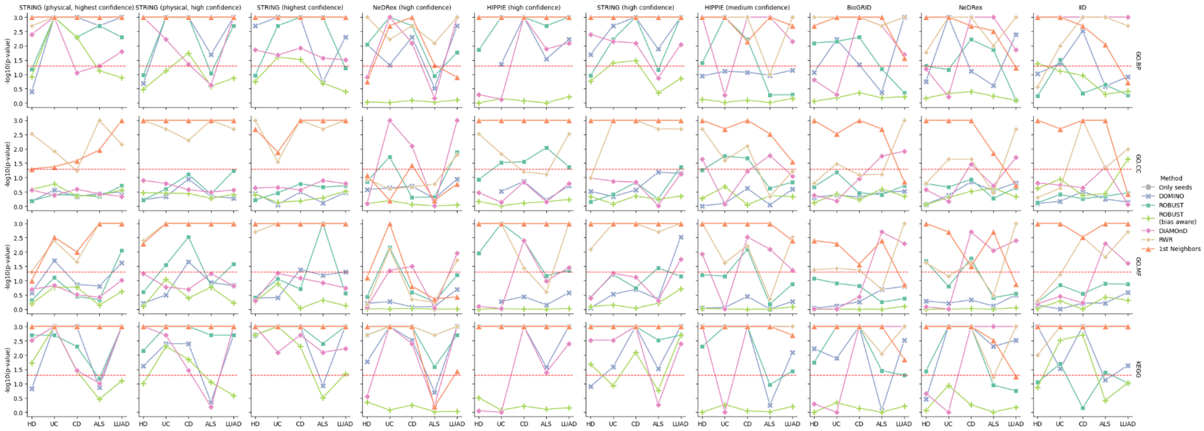


***Figure S6:*** *Results of the functional coherence analysis with DIGEST in reference-based mode for different AMIM, network, gene set source (GO.BP, GO.CC, GO.MF, or KEGG), and disease combinations. Functional coherence is expressed through empirical p-values transformed using* $-\log_{10}$. *The significance level of 0.05 is indicated by the dashed line.*
